# Three classes of epigenomic regulators converge to hyperactivate the essential maternal gene *deadhead* within a heterochromatin mini-domain

**DOI:** 10.1101/2021.05.24.445410

**Authors:** Daniela Torres-Campana, Béatrice Horard, Sandrine Denaud, Gérard Benoit, Benjamin Loppin, Guillermo A. Orsi

## Abstract

The formation of a diploid zygote is a highly complex cellular process that is entirely controlled by maternal gene products stored in the egg cytoplasm. This highly specialized transcriptional program is tightly controlled at the chromatin level in the female germline. As an extreme case in point, the massive and specific ovarian expression of the essential thioredoxin Deadhead (DHD) is critically regulated in *Drosophila* by the histone demethylase Lid and its partner, the histone deacetylase complex scaffold Sin3A, via yet unknown mechanisms. Here, we identified the Brahma chromatin remodeler sub-unit Snr1 and the insulator component Mod(mdg4) as essential for *dhd* expression and investigated how these epigenomic effectors act with Lid and Sin3A to hyperactivate *dhd*. Using Cut&Run chromatin profiling with a dedicated data analysis procedure, we found that *dhd* is intriguingly embedded in an H3K27me3/H3K9me3-enriched mini-domain flanked by DNA regulatory elements, including a *dhd* promoter-proximal element essential for its expression. Surprisingly, Lid, Sin3A, Snr1 and Mod(mdg4) impact H3K27me3 and this regulatory element in distinct manners. However, we show that these effectors activate *dhd* independently of H3K27me3/H3K9me3, and that these marks are not required to repress *dhd*. Together, our study demonstrates an atypical and critical role for chromatin regulators Lid, Sin3A, Snr1 and Mod(mdg4) to trigger tissue-specific hyperactivation within a unique heterochromatin mini-domain.

**AUTHOR SUMMARY:** Gene expression is tightly regulated by conserved protein complexes that act at the chromatin level to allow or restrict transcription. Such epigenetic control of gene activity defines the identity of different cell types during development, as well as their response to environmental cues. Yet, how multiple chromatin factors converge to achieve precise gene regulation remains difficult to address, partly due to the lack of biological situations where these intricate relationships can be studied. In this paper, we have addressed this issue by dissecting the regulation of *deadhead*, an essential gene specifically and massively expressed in the Drosophila germline. Unexpectedly, we found that its hyperactivation occurs despite *deadhead* being embedded in an apparently unfavorable chromatin mini-domain, notably featuring repressive histone modifications. We further demonstrate that four chromatin effectors, Lid, Sin3A, Snr1 and Mod(mdg4), have distinct, atypical and essential roles to ensure *deadhead* expression within this chromatin environment. Together, our findings put into perspective our understanding on these regulatory factors by illustrating how they can exert a biologically essential function via non-canonical mechanisms.

## INTRODUCTION

Gene expression is tightly controlled in eukaryotic cells by the composition, organization and dynamics of nucleosomes, consisting of an octamer of histone proteins wrapped in ∼146bp of DNA. The concerted activity of protein complexes including histone chaperones, readers and writers as well as nucleosome remodelers, defines the positioning, composition and post-translational modifications of nucleosomes [1–3]. The resulting chromatin landscape is further organized by insulator proteins that delimit tridimensional contacts along the genome, forming sub-nuclear domains and guiding contacts between promoters and their cognate regulatory elements [4]. This tightly regulated epigenomic environment profoundly influences RNA Polymerase access to DNA and transcriptional activity.

Tremendous efforts in the past decades aimed at dissecting the roles of these epigenomic effectors *in vivo*. A privileged method is ablation or dosage manipulation of each component to measure its impact on gene expression. While these approaches can yield precious functional insight, the ubiquitous expression and wide range of activities of these factors, as well as redundancies in their interactions, make it difficult to infer their precise function. Understanding their function therefore requires identifying biologically relevant situations where disrupting these effectors impacts transcription in a critical and specific manner. We previously described one of such cases, where perturbation of the histone demethylase Lid/KDM5 or the histone deacetylase scaffold Sin3A in *Drosophila* ovaries dramatically abrogated the expression of the essential maternal gene *deadhead (dhd)* [5].

The *Drosophila* egg is loaded with maternal gene products synthesized by germline nurse cells that enable early embryonic development in the absence of zygotic transcription [6]. An extreme example of this specialized transcriptome, *dhd* is among the most highly expressed genes in adult ovaries, while it is almost completely silent in any other tissue and developmental stage [5,7–9]. The DHD protein is a thioredoxin involved in regulating the general redox state in oocytes [10]. In addition, DHD plays a critical role at fertilization to reduce cysteine-cysteine disulfide bonds on the Protamine-like proteins that replace histones on chromatin during spermiogenesis [9, 11]. In the absence of DHD, paternal chromosomes fail to decondense and are excluded from the first zygotic nucleus, leading to haploid gynogenetic development and embryonic lethality. The *dhd* locus, which produces a single, short (952bp), intronless transcript is packed within a 1369bp region that separates its flanking genes *Trx-T* and *CG4198*. Remarkably, these two genes are expressed exclusively in the male germline, thereby constituting an apparently unfavorable environment for *dhd* transcription in ovaries. In addition, we showed that a 4305bp transgene spanning only *Trx-T, dhd* and part of *CG4198* largely recapitulates the expression of *dhd* [5, 9], indicating that regulatory elements sufficient for *dhd* activation are contained within this restricted region. Our previous study further found that Lid and Sin3A are essential activators of *dhd* in *Drosophila* ovaries, in striking contrast to their otherwise relatively modest impact on the rest of the transcriptome. Considering these unusual features, we postulated that the exquisite sensitivity of *dhd* to these broad-acting chromatin effectors revealed a singular mode of epigenomic regulation that enables its massive and specific ovarian expression [5].

Here, we exploited this singular model locus to understand how multiple classes of epigenomic effectors converge to achieve programmed transcriptional hyperactivation. We identified the Brahma chromatin remodeler component Snr1 [12] and the insulator complex component Mod(mdg4) [13] as factors that share with Lid and Sin3A a critical and highly specific role in activating *dhd*. By exploiting the chromatin profiling method Cut&Run [14] and an adapted data analysis strategy, we found that *dhd* is unexpectedly embedded within a heterochromatin mini-domain flanked by two border regulatory elements. One of these is a *dhd*-proximal element, which encompasses a DNA Replication-related Element (DRE-box) motif [15] that is essential for *dhd* expression. Yet, exploiting knockdown and transgenic tools, we found that Lid, Sin3A, Snr1 and Mod(mdg4) activate *dhd* independently of the associated heterochromatin mini-domain. Furthermore, this mini-domain is not required to restrict *dhd* expression to ovaries. Together, our results put into perspective our understanding on these epigenomic regulators by revealing how they exert a biologically essential control of *dhd* via non-canonical mechanisms.

## RESULTS

### Mod(mdg4) and Snr1 are essential for dhd expression

We previously performed a female germline RNA interference screen to identify chromatin factors required for paternal chromosome incorporation into the zygote at fertilization. As part of that screen, Lid and Sin3A were identified as essential regulators of *dhd* expression. Because Lid and Sin3A can interact within a co-repressor complex [16, 17], we asked whether other chromatin regulatory complexes might also be involved in *dhd* regulation. We therefore broadened our analysis to other knockdowns that caused maternal effect sterility associated with a *dhd-*like mutant phenotype, i.e. defective sperm nuclear decompaction at fertilization. Among these, we focused on two additional UAS-controlled small hairpin RNA (shRNA) constructs from the TRiP collection [18], respectively targeting *mod(mdg4)* and *Snr1*. Snr1 is an essential subunit of the Brahma chromatin remodeler that mediates protein-protein interactions within this complex as well as with external interacting partners [12, 19]. The *mod(mdg4)* gene codes for up to 31 isoforms [20], all of which are targeted by the shRNA construct. Among these, the most well characterized, Mod(mdg4)67.2 is a common component of boundary insulators in the *Drosophila* genome [21]. These two candidates belonged to two classes of epigenomic effectors distinct from Lid and Sin3A, and we thus decided to investigate their function during the oocyte to zygote transition.

When activated by the Maternal Triple Driver (MTD) Gal4 source, these shRNAs efficiently reduced the levels of *mod(mdg4)* and *Snr1* transcripts (FigS1-A,B). Previous studies reported defective oogenesis and diminished egg production in *mod(mdg4)* as well as *Snr1* mutant females [19, 22]. Consistently, females with ovarian knockdown of *mod(mdg4)* or *Snr1* (hereby referred to as *mod(mdg4)* KD or *Snr1* KD females) were almost completely sterile (Table 1). Indeed, while KD females were able to lay more eggs than mutants, these almost systematically failed to hatch. Focusing on paternal chromatin organization at fertilization in these embryos, we found that both *mod(mdg4)* and *Snr1* ovarian KDs led to failure of male pronucleus decondensation and persistence of its elongated morphology (Fig1-A). Concomitantly, these embryos exhibited retention of the protamine fusion Mst35Ba::GFP (ProtA::GFP) marker [23] in paternal chromosomes, as observed in *dhd* mutants [9, 11].

**Fig 1.**
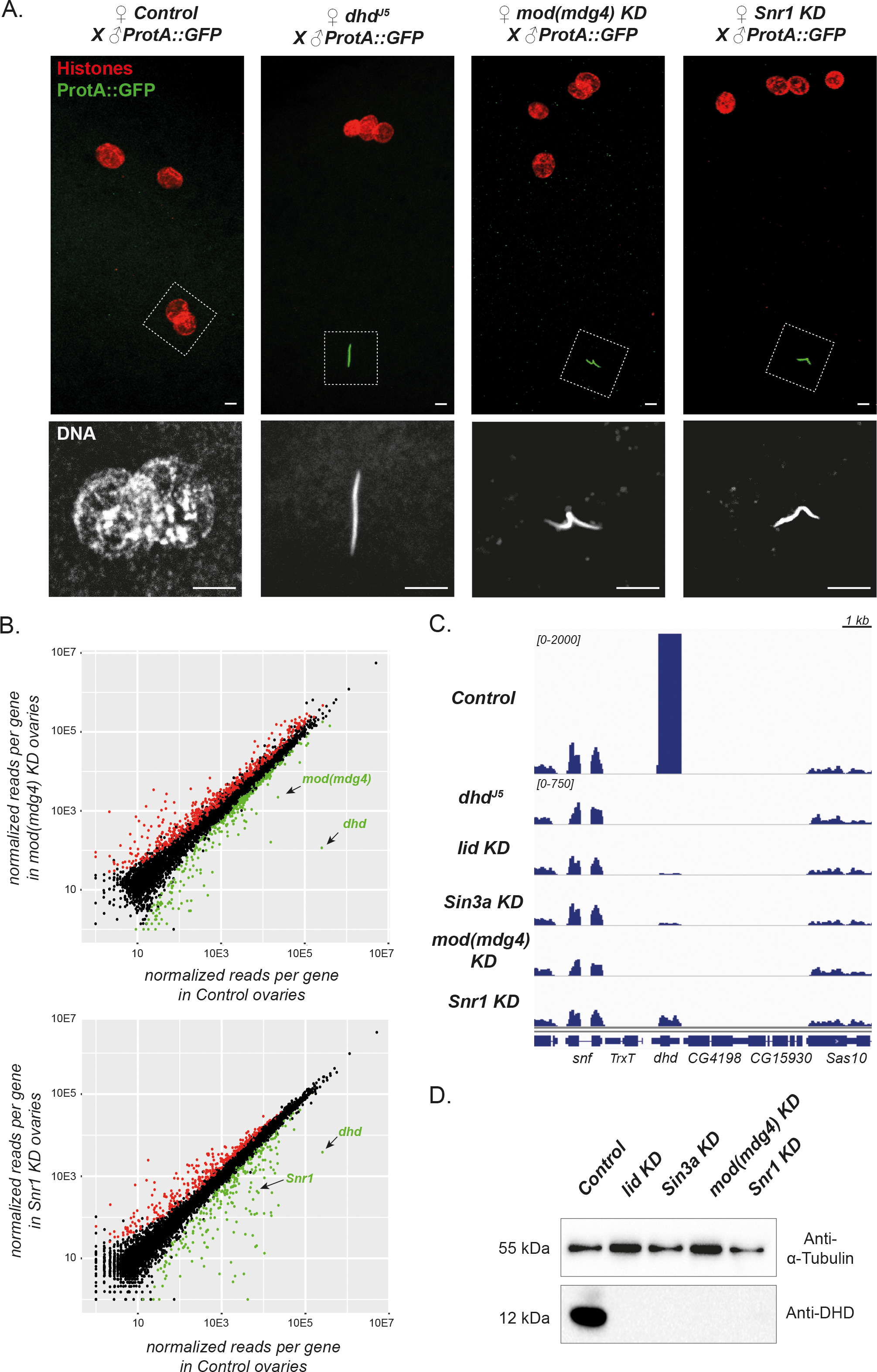
Mod(mdg4) and Snr1 are required for *dhd* expression. A—Maternal Mod(mdg4) and Snr1 are required for protamine removal and sperm nuclear decompaction at fertilization. Top: Confocal images of pronuclear apposition in eggs from Control (*MTD>+*), *dhd^J5^*, *mod(mdg4)* KD or *Snr1* KD females mated with transgenic *ProtA::GFP* males. The sperm nucleus in *dhd^J5^*, *mod(mdg4)* KD and *Snr1* KD eggs retains ProtA::GFP (green) and has a needle-shape morphology. Bottom: zoom on the sperm nucleus. Scale bars: 5μm. B— *dhd* is strongly downregulated in *mod(mdg4)* KD and *Snr1* KD ovaries. RNA-seq normalized reads per gene (in RPKM) are shown for *mod(mdg4)* KD vs Control (top) and *Snr1* KD vs Control (bottom). Genes downregulated (green) or upregulated (red) in KD ovaries are highlighted. C— Genome Browser view of Control, *dhd^J5^*, *lid* KD, *Sin3a* KD, *mod(mdg4)* KD and *Snr1* KD ovarian RNA-seq signal at the *dhd* region showing dramatic downregulation in all KD conditions. Note that the scale bar in Control ovaries is truncated for readability. D— The DHD protein is undetectable in KD ovaries. Western blot analysis using an anti-DHD antibody on ovary extracts of the indicated genotypes. Alpha-tubulin is used as a loading control.

**Table 1.**
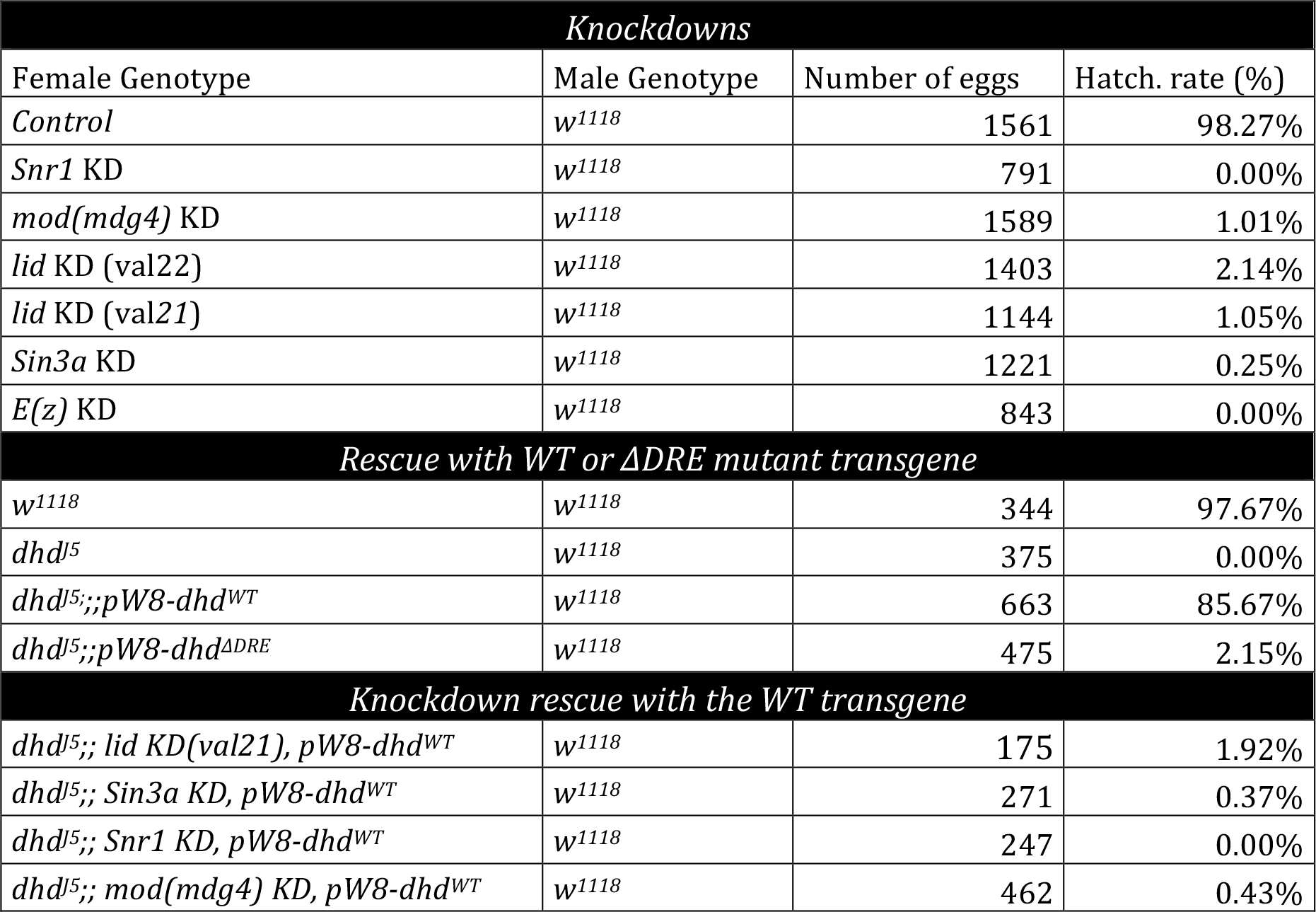
Embryo hatching rates. The *^w1118^* strain is used as reference.

The above results suggest that Mod(mdg4) and Snr1 could regulate *dhd* expression. RNA-sequencing on *mod(mdg4)* and *Snr1* KD ovaries indeed revealed that *dhd* is dramatically downregulated in both KDs, with a fold reduction of almost two orders of magnitude (Fig1-B,C, FigS1-B,E). *dhd* was the first most strongly affected gene in *mod(mdg4)* KD ovaries in terms of fold-change in expression, and the 14^th^ most affected gene in *Snr1* KD ovaries, contrasting with a more modest impact of both KDs on the rest of the transcriptome and the limited overlap in their effects (Fig S1-B,C,D). Consistently, DHD protein levels assessed by Western Blot in KD ovaries were also dramatically reduced (Fig1-D). Therefore, despite the packed genomic organization of the *dhd* locus, its expression strictly and singularly depends on multiple epigenomic effectors belonging to three distinct classes, namely histone modifiers (Lid, Sin3A), nucleosome remodelers (Snr1) and insulators (Mod(mdg4)).

### dhd lies within an H3K27me3 mini-domain flanked by DNA regulatory elements

We previously showed that H3K4me3, a histone modification associated with active transcription, is enriched at the *dhd* promoter and that this mark is lost in *lid* KD ovaries [5](FigS2-A). Further exploiting published ChIP-seq datasets, we surprisingly found that *dhd* lies within a ∼5kbp mini-domain featuring two types of repressive histone modifications: H3K27me3 and H3K9me3 [24, 25]. H3K27me3 is the hallmark of Polycomb-based repression [26, 27], whereas H3K9me3 dictates Heterochromatin Protein 1 (HP1)-based repression [28, 29]. Importantly, H3K27me3 was enriched in both somatic follicle cells and germline nurse cells (FigS2-A). This appeared at odds with the massive germline expression of *dhd*.

To more precisely characterize the *dhd* H3K27me3-enriched mini-domain, we next implemented the Cut&Run epigenomic profiling method [14]. In Cut&Run, histone modifications of interest are targeted *in situ* by a specific antibody following tissue permeabilization. Target-bound antibodies are subsequently coupled to a fusion between the bacterial Protein A and Micrococcal Nuclease (ProteinA-MNase) that cleaves exposed DNA in the vicinity of the antibody, releasing target nucleosomal particles into solution. Importantly, DNA bound by other proteins such as polymerases or DNA sequence-specific transcription factors in the immediate vicinity of the nucleosome-bound antibody is also expected to be cleaved and released (Fig2-A). In particular, DNA regulatory elements occupied by sequence-specific transcription factors are typically associated with MNase footprints distinctly shorter than nucleosomes [30–32]. Partially unwrapped dynamic nucleosomes typically associated with regulatory elements can also produce such distinctly short footprints [33]. A Cut&Run experiment should thus identify DNA regulatory elements that are in physical proximity of target histone modifications.

**Fig 2.**
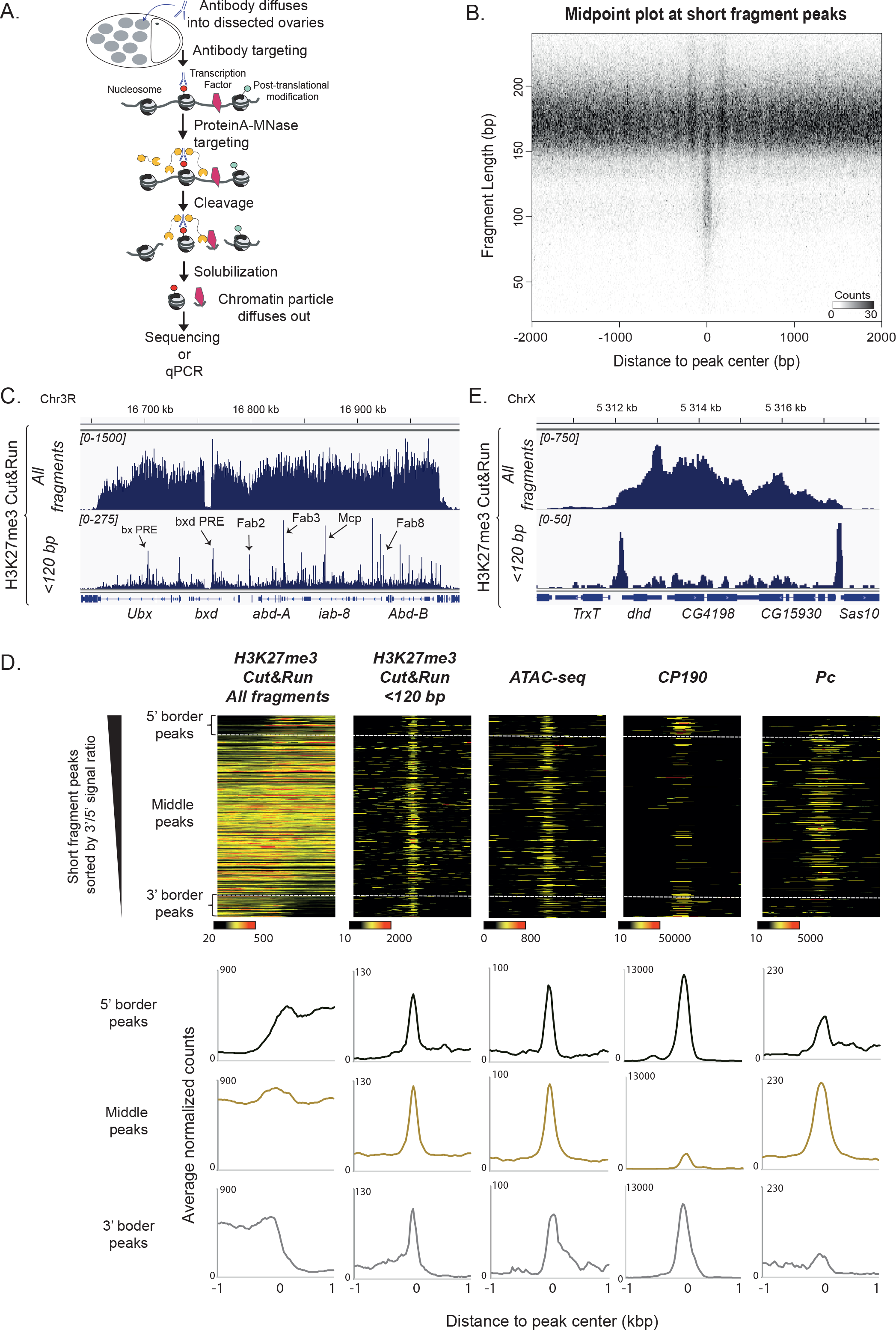
*dhd* is embedded in an H3K27me3 mini-domain flanked by regulatory elements. A— Schematic overview of the Cut&Run procedure for dissected *Drosophila* ovaries. After tissue permeabilization and antibody targeting, ProteinA-MNase cleaves nearby exposed DNA allowing the solubilization and retrieval of both nucleosomal particles carrying the targeted histone modification and DNA particles occupied by transcription factors in the immediate vicinity. B—Cut&Run reveals nucleosomes and transcription factor binding sites. Mid-point plot of ovarian H3K27me3 Cut&Run data centered at peaks identified by MACS2 from short fragments (<120bp) in the same experiment. This plot represents all paired-end sequenced fragments as their middle point coordinate in the X-axis, and their size in the Y-axis, revealing a class of clustered short fragments (50-130bp) flanked by nucleosome-sized fragments (>140bp). C— H3K27me3 Cut&Run at the bithorax complex (BX-C) in *Drosophila* ovaries reveals its regulatory architecture. Genome browser track displaying all Cut&Run fragments and <120 bp fragments separately. Multiple well-described Polycomb Response Elements (PRE) and insulators within the Bithorax complex detected as short fragment peaks are indicated (arrows). D— Cut&Run re-discovers regulatory elements associated with H3K27me3 genome-wide. Upper panels: short fragment peaks read density heatmaps of ovarian H3K27me3 Cut&Run (all fragments and <120bp fragments), ATAC-seq (from S2 cells, *Jain et al., 2020*), CP190 ChIP-seq (from Kc cells, *Li et al., 2015*) and Polycomb ChIP-seq (Pc, from S2 cells, *Enderle et al., 2011*) plotted at ±1kb around peak summit. Data is sorted by the ratio of H3K27me3 Cut&Run total reads at the 3’ versus 5’ flanks to reveal short fragment peaks at the borders or within H3K27me3 domains (dashed lines). Lower panels: average profiles corresponding to the top heatmaps, distinguishing 5’ border peaks, 3’ border peaks and peaks embedded within domains. Cut&Run short fragment peaks are enriched for ATAC-seq signal as well as CP190 (particularly at border peaks) and Polycomb (particularly at middle peaks). E— The *dhd* region features an H3K27me3 mini-domain. Genome browser snapshots showing the distribution of all fragments and <120 bp fragments in the *dhd* region, revealing that *dhd* lies within a ∼5450bp H3K27me3 mini-domain flanked by border regulatory elements.

With this in mind, we conducted H3K27me3 Cut&Run in *Drosophila* ovaries. Using only 12 pairs of ovaries per sample, we robustly revealed H3K27me3 domains. Remarkably, visualization of Cut&Run fragments shorter than 120bp (which excludes fully wrapped octameric nucleosomes) revealed that these were enriched at discrete peaks within H3K27me3 domains. Genome-wide analysis identified 679 peaks of fragments <120bp (hereon referred to as “short fragment peaks”) that were ∼250bp-wide in average (Fig2-B,D). We hypothesized that short fragment peaks represented H3K27me3-associated regulatory elements occupied by transcription factors. Within H3K27me3 domains, we expected these to include Polycomb Response Elements (PREs) as well as insulators. For example, short fragment peaks corresponded to several well-described PREs and insulators in the Bithorax complex H3K27me3 domain [34–36] (Fig2-C), consistent with observations in larval tissue [37]. To ask whether this reflects a broader genome-wide trend, we compared short fragment peaks with PRE and insulator markers genome-wide. Although there is scarce genome-wide data available for *Drosophila* ovaries, H3K27me3 domains are generally present in most cell types. We thus exploited datasets from embryonic-derived S2 and Kc cell lines. Genome-wide, small fragment peaks identified in ovaries were enriched for ATAC-seq signal [38] -revealing hyper-accessible DNA-, arguing that these indeed correspond to DNA regulatory elements (Fig2-D). Enrichment at these peaks of the Polycomb protein [39] and the insulator protein CP190 [40] further argues that these elements often correspond to functional PREs or insulators. Accordingly, at the borders of H3K27me3 domains, short fragment peaks were more frequently associated with CP190, confirming previous reports that this factor is associated with H3K27me3 domain boundaries [21, 41] (Fig2-D). Instead, Polycomb was rather enriched at peaks localized internally within these domains. Our Cut&Run analysis strategy therefore revealed not only the breadth of H3K27me3 domains in ovaries but also their associated DNA regulatory elements.

With this approach, we next confirmed that *dhd* is included in a ∼5450bp H3K27me3 mini-domain that extends from the promoter region of *dhd* to the promoter of the next gene active in ovaries, *Sas10*. Surprisingly, short fragment analysis revealed two DNA regulatory elements associated with H3K27me3, precisely at the mini-domain borders, with no internal peaks present (Fig2-E). This regulatory architecture was quite unusual, as we could not find any other H3K27me3 domain in the genome sharing this particular organization with two border elements and no internal elements. ChIP-seq data from S2 and Kc cells further confirmed that this mini-domain is present in both cell lines, with enrichment in ATAC-seq signal at the border elements (FigS2-B). In addition, in Kc cells, *dhd* border elements are occupied by CP190 and Mod(mdg4), both of which can be found at the boundaries of *Drosophila* H3K27me3 domains [21, 42]. Finally, the *dhd*-proximal 5’ border element featured a significant, although very modest enrichment for PRE markers Polycomb and Polyhomeotic (Fig-S2-B). Together, these results revealed that *dhd* lies within a unique H3K27me3 mini-domain featuring only border DNA regulatory elements.

### Sin3A, Snr1 and Mod(mdg4) control the regulatory architecture of the dhd mini-domain

Previous reports showed that depletion of insulator proteins Mod(mdg4), as well as CTCF, Su(Hw), CP190 or BEAF-32, did not affect the spread of Polycomb-associated domains but instead caused a general decrease in H3K27me3 levels [21]. Therefore, Mod(mdg4) was expected to act as a transcriptional repressor at H3K27me3 domains. Yet, this is in contrast to the negative effect of *mod(mdg4)* KD on *dhd* expression. Furthermore, the genome-wide impact of Lid, Sin3A or Snr1 on H3K27me3 in *Drosophila* has not been evaluated. To evaluate the potential role for all these factors in regulating the *dhd* H3K27me3 mini-domain, we analyzed this mark in KD ovaries. As a control, we included a KD for the H3K27 methyltransferase Enhancer of zeste (E(z)). While *E(z)* KD females were sterile as previously described [43, 44] (Table1), they were able to lay eggs and displayed only a moderate effect on *dhd* expression (a 25% reduction compared to controls) (FigS3-A). Immunofluorescence staining on whole dissected control ovaries showed that H3K27me3 marks follicle cell nuclei, the karyosome (i.e the oocyte nucleus) and nurse cell nuclei, although nurse cell staining was relatively weaker (FigS3-B), consistent with previous reports [43]. As expected, H3K27me3 was undetectable in the karyosome and in nurse cells of *E(z)* KD ovaries, whereas follicle cells (which do not express MTD-driven shRNAs) still carried this mark at normal levels. While *lid*, *Sin3a* and *mod(mdg4)* KD ovaries displayed normal H3K27me3 staining, we observed a moderate reduction in H3K27me3 levels in nurse cells in *Snr1* KD ovaries, even while H3K27me3 levels were not affected on the karyosome (FigS3-B).

We next carried out H3K27me3 Cut&Run on ovaries from all KDs. We first segmented the Cut&Run H3K27me3 signal in control ovaries to identify 278 H3K27me3 domains, ranging from 3 to 240kb in width. Within these domains, we compared the average enrichment in H3K27me3 signal in control and KD ovaries (Fig3-A). In *E(z)* KD ovaries, Cut&Run experiments revealed only a moderate loss of H3K27me3 signal (35% average reduction at these domains compared to controls) (Fig3-A), contrasting with the strong global reduction in H3K27me3 immunofluorescence signal. This difference is likely to reflect the fact that the H3K27me3 signal from Cut&Run experiments originates from both germline and somatic cells. Accordingly, *E(z)* KD completely abrogated H3K27me3 signal at the *spen*, *Corto* or *ptc* loci, all of which are decorated with H3K27me3 in nurse cells but not in follicle cells (FigS3-C) [24]. In contrast, the *gl, dpp,* or *repo* loci, which show stronger H3K27me3 in follicle cells compared to nurse cells, were only slightly affected in *E(z)* KD ovaries (FigS3-C). Together, these results show that our Cut&Run strategy detects H3K27me3 signal from both germline and somatic cells and is able to detect quantitative differences in the averaged signal when nurse cells are strongly affected. Consistent with immunofluorescence experiments, *lid, Sin3a* and *mod(mdg4)* KDs had only a modest global impact on average H3K27me3 levels (5% reduction compared to controls), and no effect on the spread of H3K27me3 domains (Fig3-A). Also consistent with our immunofluorescence experiments, *Snr1* KD led to a more severe average reduction of H3K27me3 Cut&Run signal compared to controls (20%), although not as dramatic as *E(z)* KD.

**Fig 3.**
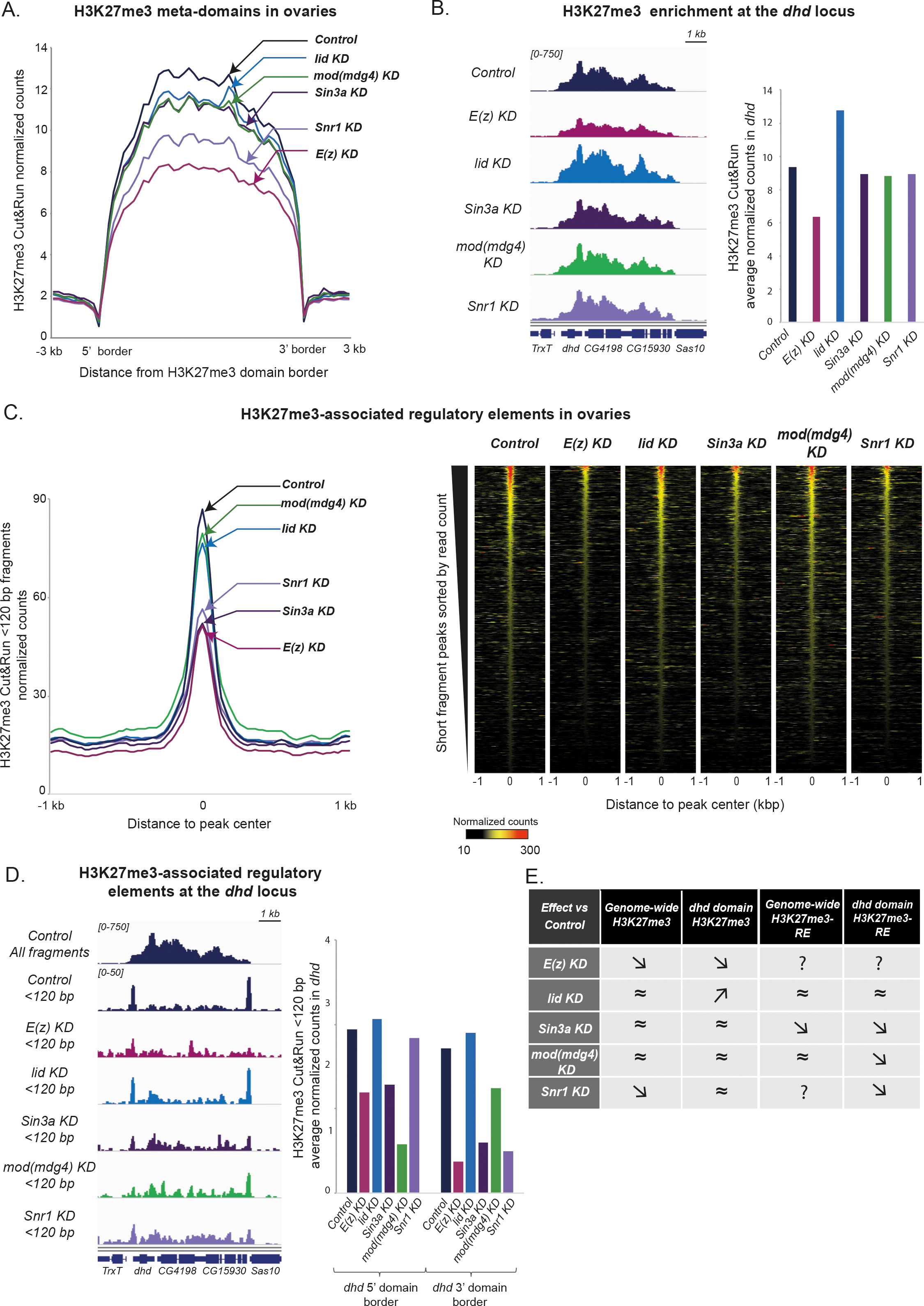
Sin3A, Snr1 and Mod(mdg4) control the regulatory architecture of the *dhd* H3K27me3 mini-domain. A– Effect of the KDs on H3K27me3 enrichment genome-wide. Average normalized counts of H3K27me3 Cut&Run (all fragments) in H3K27me3 domains (plotted as meta-domains and including ±3kb from domain borders) in Control (*MTD>+*), *lid* KD, *Sin3a* KD, *mod(mdg4)* KD, *Snr1* KD and *E(z)* KD (arrows). B– *lid* KD, but not *Sin3a*, *Snr1* or *mod(mdg4)*, impacts H3K27me3 enrichment at the *dhd* mini-domain. Left: genome browser plots of normalized H3K27me3 Cut&Run signal (all fragments) at the *dhd* genomic region in Control and KD ovaries. Right: Quantification of normalized read counts for the same samples. Replicates are shown in FigS4-A. C– *Sin3a* and *Snr1* KD, but not *lid* or *mod(mdg4)*, impact H3K27me3-associated regulatory elements genome-wide. Left: H3K27me3 Cut&Run <120 bp fragments normalized counts in Control and KD ovaries, plotted at ±1kb around the summit of short fragment peaks. Right: Heatmaps displaying H3K27me3 Cut&Run short fragment peaks normalized read counts ±1kb around peak center in Control and KD ovaries. D-Sin3A, Snr1 and Mod(mdg4), but not Lid, impact the organization of regulatory elements at the borders of the *dhd* H3K27me3 mini-domain. Left: genome browser plots of normalized Control H3K27me3 Cut&Run signal (all fragments, top) and of normalized signal from <120bp fragments retrieved in H3K27me3 Cut&Run in Control and KD ovaries. Right: Quantification of <120bp fragments normalized read counts for the same samples. 5’ and 3’ border elements are plotted separately. Replicates are shown in FigS4-C. F-Table recapitulating the effect of the different KDs on H3K27me3 and H3K27me3-associated regulatory elements (H3K27me3-RE) genome-wide and at the *dhd* locus. “≈” indicates modest or no change, “↗” indicates an increase and “↘” a decrease in average read counts compared to Control. “?” indicates inability to conclude.

In agreement with genome-wide observations, the levels of H3K27me3 in the *dhd* mini-domain were reduced in *E(z)* KD ovaries and unaffected in *Sin3a* or *mod(mdg4)* KD ovaries. More surprisingly, the domain was also unaffected in *Snr1* KD ovaries, despite the fact that H3K27me3 is globally impacted by this knockdown (Fig3-B, FigS4-A). Within the sensitivity limits of our approach, these results indicate that *Sin3a, Snr1* and *mod(mdg4)* KDs have little if any impact on H3K27me3 at the *dhd* locus. Conversely, in *lid* KD ovaries, in which global H3K27me3 levels were unaffected, we detected an increase in H3K27me3 levels at the *dhd* mini-domain (Fig3-A,B, FigS4-A). This raised the possibility that Lid could facilitate *dhd* expression by counteracting Polycomb-mediated repression.

Since the *dhd* mini-domain also featured H3K9me3, we next turned to Cut&Run followed by qPCR to evaluate its status in KD ovaries. H3K27me3 Cut&Run-qPCR confirmed that this approach quantitatively measures the expected enrichments at H3K27me3 domains and detects biologically relevant variations in the signal (FigS4-B). To validate the H3K9me3 Cut&Run-qPCR approach in ovaries, we exploited the *CG12239* gene as a positive control [25], and detected an expected enrichment in H3K9me3 signal at this locus (FigS4-B). At the *dhd* locus, H3K9me3 was enriched as expected from ChIP-seq results. Importantly, knockdown of *lid, Sin3a, mod(mdg4) or Snr1* had no effect on this enrichment (FigS4-B). The *dhd* heterochromatin mini-domain including H3K27me3 and H3K9me3 is thus independent of Sin3A, Snr1 and Mod(mdg4), whereas Lid counteracts H3K27me3.

We next evaluated the impact of different KDs on the *dhd* mini-domain short fragment peaks at border regions. We first analyzed the effect of our different knockdowns on the full set of 679 peaks previously defined (Fig2-B). Both *E(z)* and *Snr1* KD led to a strong (∼63%) decrease in short fragment peak average counts genome-wide (Fig3-C). Since these KDs also affect global H3K27me3 levels, this reduction could result from a general absence of histone modification-targeted MNase on chromatin. Remarkably, *Sin3a* KD led to a similarly strong effect on short fragment peak counts that could not be attributed to its global impact on H3K27me3. Instead, this data suggests that Sin3A is required to ensure proper occupancy and organization of transcription factors and/or nucleosomes at DNA regulatory elements associated with H3K27me3. In contrast, *lid* or *mod(mdg4)* KD did not globally affect short fragment peak counts, indicating that these factors do not play such a role (Fig3-C).

Consistent with their effects genome-wide, short fragment counts at the *dhd* mini-domain border elements were strongly diminished upon *E(z)* and *Sin3a* KDs (Fig3-D). Intriguingly, *mod(mdg4)* KD led to a similar impact on these border elements (particularly the *dhd-*proximal one), even though it did not globally affect H3K27me3-associated elements genome-wide (Fig3-D). This observation could indicate that the *dhd* border elements become less frequently occupied by transcription factors, that these factors become less frequently associated with H3K27me3, and/or that their nucleosomal organization is compromised. In all cases, this suggests that Mod(mdg4) is required to ensure chromatin organization of the border DNA regulatory elements at the *dhd* mini-domain. Remarkably, *Snr1* KD led to a similar effect on border elements without affecting H3K27me3 levels at the *dhd* mini-domain, suggesting that Snr1 is also required for the proper organization of the *dhd* border elements. In striking contrast, *lid* KD had no detectable effect on these regulatory elements (Fig3-D). We concluded that Lid, although essential for *dhd* expression, was not required to ensure the proper organization of *dhd* border elements.

Altogether our results, summarized in Fig3-E, indicate that Lid, Sin3A, Snr1 and Mod(mdg4), impact H3K27me3 or its associated regulatory elements genome-wide and/or at the *dhd* mini-domain in four distinct manners.

### The dhd promoter-proximal DNA regulatory element is required for dhd expression independently of its heterochromatin mini-domain

We next performed sequence analysis of the *dhd* mini-domain border elements, screening against the flyreg.v2 [45, 46] transcription factor DNA binding motif database. At the 5’ border element, which mapped to the *dhd* promoter region, we identified four perfect matches for the DNA replication-related element (DRE) motif, TATCGATA (Fig4-A). This motif is recognized by the insulator-associated factor BEAF-32 [47] and the core-promoter factor DREF [15]. These four DRE motifs overlap in the palindromic sequence TATCGATATCGATA, 37bp upstream of the *dhd* transcription start site. Consistently, BEAF-32 and DREF both occupy this element in Kc cells (FigS5) [40]. Previous studies showed that BEAF-32 null females are partially fertile (∼40% hatching rate) [48], indicating that this factor is not essential for *dhd* expression. In turn, DREF is essential in a cell-autonomous manner and indeed *dref* mutations cause oogenesis defects [49]. Accordingly, we observed severe atrophy and failure to produce oocytes in *dref* KD ovaries. Because this precluded studying the role of DREF in *dhd* regulation, we instead sought to probe the importance of the DRE motifs themselves.

**Fig 4.**
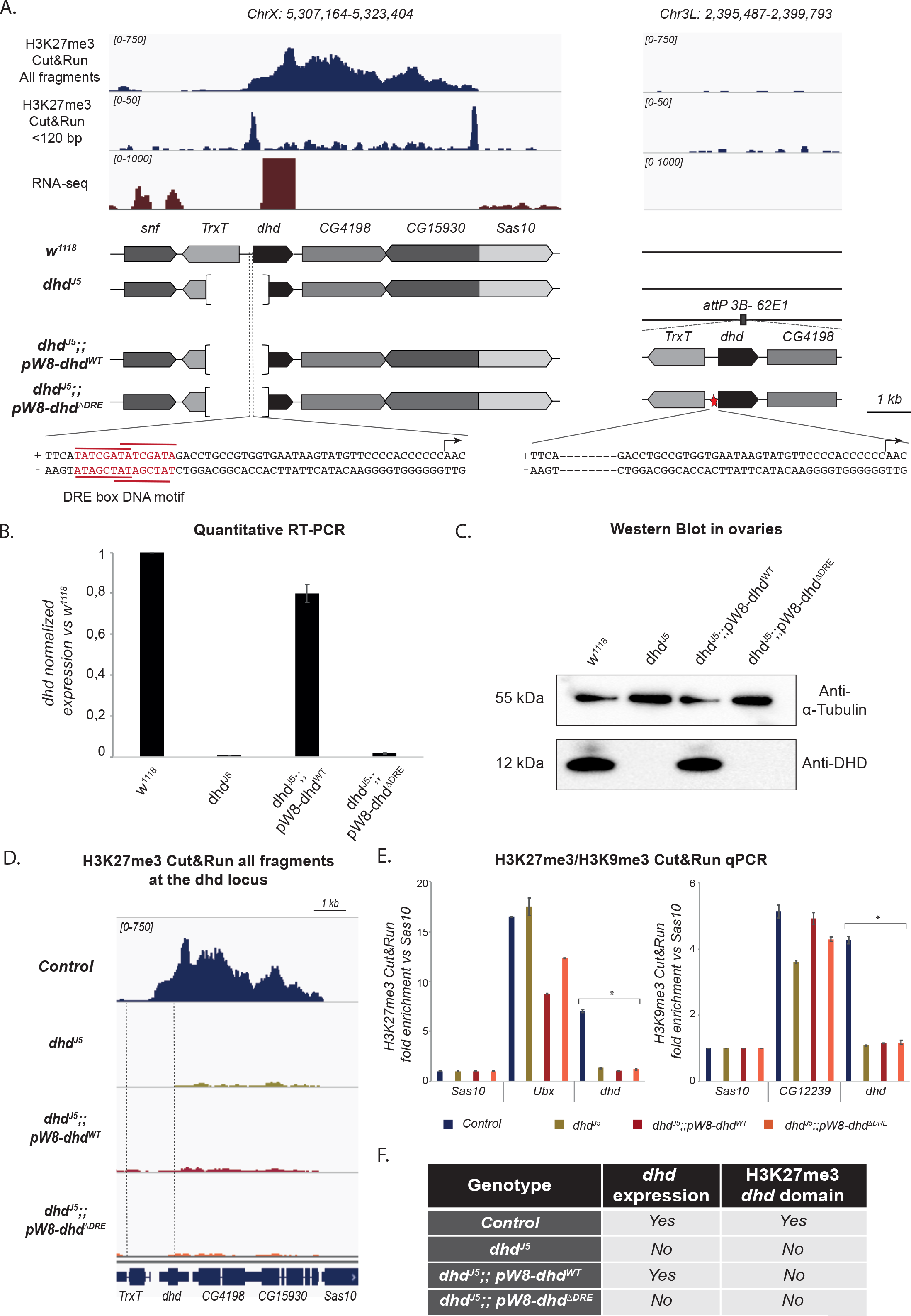
The *dhd* promoter-proximal DRE motifs are required for its expression. A– Schematic representation of the genotypes studied in this figure. Upper panels: genomic browser views recapitulating the Control distribution of H3K27me3 Cut&Run signal (all fragments and <120bp fragments) as well as RNA-seq signal from Figures 1 and 2 at the *dhd* locus and showing lack of signal at the transgene insertion locus in the absence of any transgenic construct. Middle panel: schematic representation of the genomic composition of *w^1118^* (reference strain), mutant and rescue flies, indicating the status of the *dhd* locus and the composition of the rescue transgene. Bottom panel: sequence of the *dhd* promoter at the endogenous location (left) and in the ΔDRE mutant transgene where the 14bp containing the DRE motifs were deleted. B- The DRE motifs at the *dhd* promoter are necessary for its expression. RT-qPCR quantification of *dhd* mRNA levels in ovaries from wild-type *w^1118^* flies, *dhd^J5^* mutants or *dhd^J5^* mutants carrying either a WT (*pW8-dhd^WT^*) or a mutant (*pW8-dhd^ΔDRE^*) transgene (normalized to *Rp49* and relative to expression in *w^1118^*). Data from biological duplicates analyzed in technical duplicates are presented as mean ± SEM. C- Western blot analysis of DHD expression in ovaries of indicated genotypes. Alpha-tubulin detection is used as a loading control. D- The H3K27me3 *dhd* domain is lost in *dhd*-containing transgenic constructs. Genome browser plots of normalized H3K27me3 Cut&Run signal at the *dhd* genomic region in Control, *dhd^J5^, dhd^J5^;;pW8-dhd^WT^ and dhd^J5^;;pW8-dhd^ΔDRE^* ovaries. The H3K27me3 domain is abolished in all genotypes except for Control. The dashed line covers the deleted segment in *dhd^J5^*. E- *dhd-*containing transgenic constructs do not recapitulate its heterochromatin domain. H3K27me3 and H3K9me3 Cut&Run-qPCR in the same genotypes as in D. The *Sas10* gene was used as a negative control and positive controls were *Ubx* for H3K27me3 and *CG12239* for H3K9me3. Fold enrichment was calculated relative to *Sas10.* Error bars show technical variability from a representative replicate. *: p-value<0.0001 in one-way ANOVA with Dunett’s multiple comparisons test to a control. F- Table summarizing the results on *dhd* expression and on the presence of *dhd* H3K27me3 mini-domain in the indicated genotypes.

The *dhd^J5^* null allele is a 1.4 kb deletion affecting the entire promoter region including the promoter-proximal regulatory element, and part of the coding region of *dhd* [7, 9] (Fig4-A). A *pW8-dhd^WT^* transgenic construct, bearing the entire *dhd* gene -including its promoter region-, restores *dhd* expression as well as fertility in *dhd^J5^* mutants [9] (Fig4-A,B, Table1). We now constructed a second rescue transgene based on the *pW8-dhd^WT^*, where the 14bp carrying the DRE motifs were deleted (*pW8-dhd^ΔDRE^*) (Fig4-A). These constructs were inserted into the same genomic location as *pW8-dhd^WT^* (62E1) and combined with the *dhd^J5^* deficiency. In striking contrast to *pW8-dhd^WT^*, the *pW8-dhd^ΔDRE^* construct was unable to restore DHD protein levels, rescue *dhd* expression, or substantially improve fertility in *dhd^J5^* deficient flies (Fig4-B,C, Table1). The DRE motifs are thus essential to ensure *dhd* expression.

To test a role for this regulatory element and its DRE motifs in regulating the H3K27me3/H3K9me3 mini-domain, we performed Cut&Run-seq and Cut&Run-qPCR on homozygous *dhd^J5^* ovaries, as well as rescue *dhd^J5^;;pW8-dhd^WT^* and non-rescued *dhd^J5^;;pW8-dhd^ΔDRE^* ovaries. Strikingly, the 5.4kbp *dhd* H3K27me3 mini-domain was completely lost in *dhd^J5^* ovaries (Fig4-D,E), despite the fact that 90% of this domain were intact in the deficient chromosome. This indicates that the *dhd-*proximal border of this mini-domain is essential for establishment and/or maintenance of H3K27me3. Furthermore, H3K27me3 signal was absent within the mini-domain in *dhd^J5^;;pW8-dhd^WT^* rescue ovaries (Fig4-D,E), suggesting that the 5’-most 2.8kbp of the domain are also insufficient to establish and/or maintain H3K27me3. This result further confirms that *dhd* can be expressed at high levels in the absence of H3K27me3, consistent with results from *E(z)* KD ovaries (FigS3-A). Finally, the H3K27me3 mini-domain was also completely absent in *dhd^J5^;;pW8-dhd^ΔDRE^* ovaries (Fig4-D,E), indicating that the DRE motifs are required for *dhd* expression independently of H3K27me3. Importantly, we also found that H3K9me3, measured by Cut&Run-qPCR, was absent from *dhd* in *dhd^J5^* as well as *dhd^J5^;;pW8-dhd^WT^* and *dhd^J5^;;pW8-dhd^ΔDRE^* ovaries (Fig4-E). Together, these results indicate that the boundary regions of the *dhd* mini-domain are individually insufficient to establish H3K27me3 or H3K9me3, and that *dhd* expression can proceed at almost normal levels independently of these marks (Fig4-F).

### Lid, Sin3A, Snr1 and Mod(mdg4) activate dhd independently of its heterochromatin mini-domain

The fact that the *pW8-dhd^WT^* transgene restored most of *dhd* expression without re-establishment of the heterochromatin mini-domain at this locus provided an opportunity to clarify the role of our set of *dhd* regulators. KD of lid is associated with increased H3K27me3 at the *dhd* mini-domain, suggesting that Lid may operate as an anti-repressor by counteracting heterochromatinization of the locus. However, we have previously found that *dhd* expression is not re-established in *lid* KD ovaries carrying a *pW8-dhd^WT^* rescue transgene [5]. Lid is thus required for *dhd* expression not only at its endogenous locus but also from the rescue transgene not decorated by H3K27me3 (Fig4). Therefore, Lid activates dhd independently of heterochromatin, suggesting that it does not operate strictly as an anti-repressor.

To discriminate between anti-repressive or activating roles of Sin3A, Mod(msg4) and Snr1, we generated flies combining a *dhd^J5^* deficiency, the *pW8-dhd^WT^* transgene and an shRNA targeting *lid*, *Sin3a, Snr1 or mod(mdg4)*, driven in germ cells by a *nos-Gal4* driver (Fig5-A). We confirmed by RT-qPCR that knockdowns were still efficient when using this driver (Fig-S6). Remarkably, all of these flies were almost completely sterile, and showed strong downregulation of *dhd* revealed by RT-qPCR (Fig5-B, Table1). Using Cut&Run-qPCR at the *dhd* locus we further confirmed that these knockdowns had no effect on H3K27me3, which remained depleted in all conditions (Fig5-C). Lid, Sin3A, Snr1 and Mod(mdg4) therefore activate *dhd* independently of its heterochromatin mini-domain.

**Fig 5.**
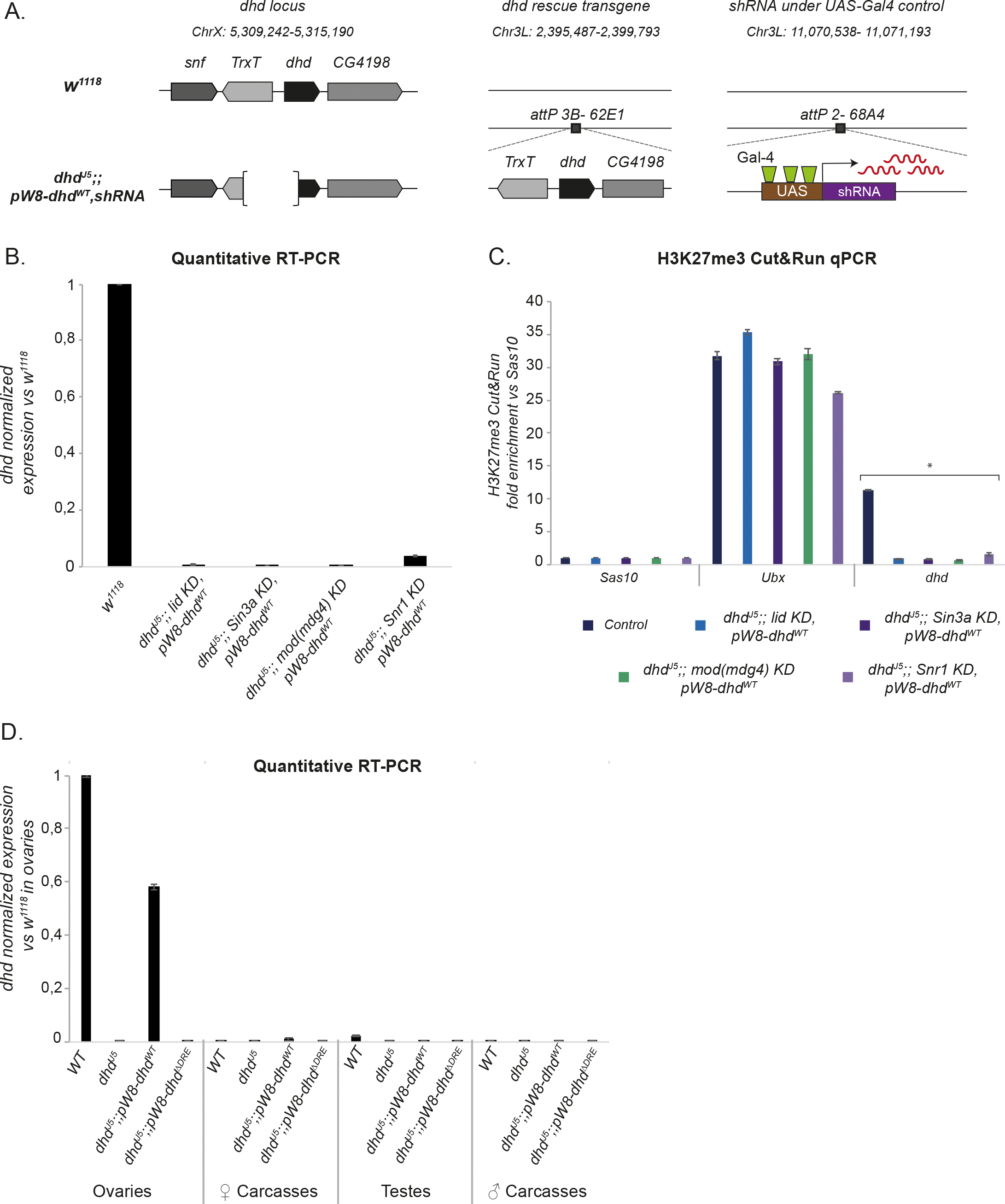
Lid, Sin3A, Mod(mdg4) and Snr1 are necessary for *dhd* expression in the absence of its heterochromatin domain. A–Schematic representation of the genomic composition of *w^1118^* (reference strain) and mutant flies carrying a rescue transgene and shRNA constructs controlled by the female germline specific nanos-Gal4 driver, respectively inserted at the platforms *attP 3B-62E1* and *attP 2-68A4*. B-The rescue transgene does not restore *dhd* expression in KD ovaries. RT-qPCR quantification of *dhd* mRNA levels in ovaries of the indicated genotypes (normalized to *Rp49* and relative to expression in *w^1118^* ovaries). Data from biological duplicates analyzed in technical duplicates are presented as mean ± SEM. C-H3K27me3 is absent from the *dhd* rescue transgene. H3K27me3 Cut&Run-qPCR in the indicated genotypes. The *Sas10* gene was used as negative control and *Ubx* as positive control. Fold enrichment was calculated relative to *Sas10.* Error bars show technical variability from a representative replicate. *: p-value<0.0001 in one-way ANOVA with Dunett’s multiple comparisons test to a control. D- *dhd* is not ectopically expressed in adult tissues in the absence of its heterochromatin domain. RT-qPCR quantification of *dhd* mRNA levels in dissected ovaries or corresponding female carcasses as well as testes or corresponding male carcasses, in all indicated genotypes (normalized to *Rp49* and relative to expression in ovaries in *w^1118^*). Data from biological duplicates analyzed in technical duplicates are presented as mean ± SEM.

### The dhd heterochromatin mini-domain is not required for dhd silencing in adult flies

Our results suggest that the *dhd* heterochromatin mini-domain does not play a repressive role in ovaries, but do not exclude that it might be required to maintain *dhd* silent in other tissues. RT-qPCR analysis on dissected ovaries, testes and male and female carcasses from transgenic lines expressing *dhd^J5^*;;*pW8-dhd^WT^* revealed *dhd* expression uniquely from ovaries (Fig5-D). Because this transgene rescues *dhd* expression without restoring heterochromatin marks, we concluded that the *dhd* heterochromatin mini-domain is not essential to repress ectopic *dhd* expression in adults, although we cannot exclude that *dhd* was weakly and/or transiently expressed in certain cell types in these conditions.

## DISCUSSION

### The ovarian hyperactivation of *dhd*

Here, we sought to understand how the genomic and epigenomic environments of *dhd* contributed to its remarkable regulation, its expression being both among the highest in Drosophila, and absolutely specific to adult ovaries [5, 9]. While Lid, Sin3A, Snr1 and Mod(mdg4) all shared a critical role in ensuring *dhd* expression, *dhd* was by far the most strongly dependent on these factors. Yet, these four broadly expressed proteins play multiple roles other than *dhd* regulation. For example, transcriptomic analyses following individual depletion of Lid, Sin3A or Snr1 in S2 cells, wing discs or pupae leads to activation or repression of hundreds of targets [16,19,50]. ChIP-seq data further indicates that Mod(mdg4), Sin3A and Lid each target several thousand sites in the genome [21,36,51,52]. Consistently, our RNA-seq analyses did reveal that each of these knockdowns were associated to up- or down-regulation of 407 to 2020 genes in ovaries. The fact that knockout of these genes typically leads to more severe phenotypic defects further suggests that their dose reduction in knockdown conditions has only limited effects, with the singular exception of *dhd*. We propose that *dhd* is a hypersensitive gene that reacts to global imbalances in the epigenome far more radically than any other locus.

The key question is therefore what is the formula for *dhd* ovarian hyperactivation. One reasonable hypothesis was that *dhd* could be highly regulated by distal enhancers. This would be notably consistent with the previously described role of Mod(mdg4) in organizing 3D contacts between regulatory elements and promoters [36]. It would also be consistent with recent findings that H3K27me3 micro-domains may reflect such contacts [53]. However, no interaction between *dhd* and any other locus can be found in Hi-C data, and our rescue transgene experiments show that a small (4kb), ectopic genomic segment encapsulates the elements required for ovary-specific hyperactivation, arguing against a critical role of the genomic and epigenomic environment in the *dhd* locus.

We indeed found a key regulatory element containing a tandem DRE motif, known to recruit the DREF core promoter factor. The minimal DRE motif (TATCGATA) is found in thousands of gene promoters [54], while multiple genes were individually shown to require this motif for proper activation. These include genes with ovarian expression, and, accordingly, DREF mutations cause oogenesis defects and female sterility [49]. In contrast, the particular tandem DRE motif in the *dhd* regulatory sequence is uncommon, being only found in 9 other gene promoters. Yet, among these 9 genes, only 4 displayed an expression bias in ovaries, and none were nearly as highly transcribed as *dhd*. Therefore, this motif does not seem to be autonomously sufficient for ovarian hyperexpression.

Another unusual feature of *dhd* is its surrounding heterochromatin mini-domain bearing both H3K27me3 and H3K9me3 marks, as well as H3K4me3. The co-occurrence of these active and repressive modifications at an ensemble of developmentally regulated genes in mammals led to the concept of bivalent promoters [55]. It is speculated that such promoters may be poised for rapid activation or repression upon differentiation. In *Drosophila*, bivalent chromatin is associated with genes that can be strongly activated in a tissue-specific manner [56, 57]. Our experiments showed that *dhd* is expressed at ∼60-70% of its normal levels in *E(z)* KD ovaries, as well as in rescue transgenes -both conditions in which the H3K27me3 mini-domain is impaired. We thus cannot exclude that H3K27me3 plays a positive role in *dhd* activation to ensure its transcription at maximum capacity, perhaps via establishment of a bivalent configuration.

Altogether, we uncovered multiple unusual genomic and epigenomic characteristics at the *dhd* locus, but failed to identify any single feature that was truly defining. The dramatic regulation of *dhd* may rely not on any individual trait but rather on a unique combination of such rare features. Further work will be needed to elucidate how these different components may together achieve ovarian hyperexpression.

### A non-canonical chromatin domain

The unique properties of *dhd* led us to uncover interesting features of its epigenomic regulators. First, *dhd* is embedded in an H3K27me3 mini-domain flanked by regulatory elements. A recent report suggested that H3K27me3 domain borders may be established independently of PREs or border elements, provided that an immediately neighboring active gene instead delimits H3K27me3 spreading [42]. The case of *dhd* is however peculiar in that the H3K27me3 domain border overlaps with this highly active gene, a scenario that was not found in other domains.

Another recent study reported the existence of H3K27me3 micro-domains (typically 2-8 nucleosomes wide) that depend on 3D contacts with larger H3K27me3 domains, mediated, in particular, by BEAF-32 and CP190 [53]. The *dhd* mini-domain is wider and much more strongly enriched in H3K27me3 than typical micro-domains. Nonetheless, our data is consistent with a model whereby H3K27me3 could be deposited via such looping interactions. First, BEAF-32 and CP190 are indeed found at the border elements of the *dhd* mini-domain. Second, this mini-domain does not feature internal PREs, arguing against an autonomous recruitment of E(z). Finally, a deletion of the BEAF-32/CP190-bearing regulatory element in the *dhd^J5^* mutant, or its displacement to an ectopic genomic location in the *pW8-dhd^WT^* transgene both abrogate H3K27me3 deposition. Consistent with such a model, data from Heurteau *et al.* show a modest reduction of H3K27me3 enrichment at the *dhd* mini-domain upon BEAF-32 depletion. Of note, BEAF-32 was also previously shown to facilitate H3K9me3 deposition at sites featuring multiple instances of the CGATA motif, alike those found at the *dhd* promoter [58]. Other studies found that ATCGAT motifs recognized by BEAF-32, also found at the *dhd* promoter, are more broadly enriched at the promoters of Lid-activated genes [59], which is the case of *dhd*. Thus, it is possible that a BEAF-32-mediated looping mechanism is responsible for H3K27me3 enrichment at the *dhd* mini-domain. However, our results also show that this mark is not strictly required to repress nor to activate *dhd* in adults, and that Lid, Sin3A, Snr1 and Mod(mdg4) activate *dhd* independently of it.

Scrutiny of *dhd* regulation further uncovered how its four regulators have convergent yet distinct roles. This was particularly intriguing for Lid and Sin3A, which can be found in a co-repressor complex [17], at odds with their positive impact on *dhd*. Indeed, their dual depletion in cultured cells causes the misregulation of hundreds of genes [16]. Interestingly, in that study, only 55 out of 849 affected genes were similarly impacted by individual and dual knockdowns, indicating that Lid and Sin3A functionally cooperate only at a minor subset of their common targets. This seems to be the case at the *dhd* locus, where individual KD of these factors caused an equally catastrophic collapse of transcriptional activity, suggesting a cooperative activity. Yet, Lid, but not Sin3A, acted as a negative regulator of H3K27me3 at the *dhd* locus, revealing at least partially independent functions. In contrast, Sin3A, but not Lid, controlled the stability of regulatory elements associated with this H3K27me3, not only at the *dhd* domain but also genome-wide.

Our results further show a critical role for Mod(mdg4) as a direct activator. In cell lines, ChIP-seq experiments specifically mapping the insulating Mod(mdg4)67.2 isoform or total Mod(mdg4) showed that additional isoforms are recruited to DNA [21]. Isoforms other than the 67.2 were found in particular at gene promoters in ovaries and female heads [4]. Such is the case at the *dhd* promoter, where total Mod(mdg4) is found but not the 67.2 isoform (FigS2-B). Non-insulating roles of Mod(mdg4) were previously discussed in the context of the Polycomb-repressed Bithorax complex where the close binding of Mod(mdg4) to *Abd-B* transcription start sites suggested a role in transcription activation [36]. In agreement, Mod(mdg4) appears to be essential to activate *dhd* within its H3K27me3 mini-domain, seemingly by stabilizing the *dhd* promoter regulatory element, although its function is equally essential in the absence of heterochromatin in the *dhd* transgenic rescue construct.

The Snr1-containing Brahma complex is required for activation of target genes in *Drosophila in vivo*, notably during immune responses [60] and tissue regeneration [50]. In ovaries, while Snr1 has a global impact on nuclear integrity and architecture, previous immunostaining experiments interestingly showed that this factor is only expressed during a restricted time in early oogenesis [19]. This underlines the fact that *dhd* may be dynamically regulated during oogenesis, with different regulatory components intervening at particular times. Considering that *Snr1* KD causes a disruption of the *dhd* promoter-proximal regulatory element associated with H3K27me3, this would suggest that its associated DNA-binding transcription factors also intervene during a restricted time in oogenesis. A precise dissection of the timing of *dhd* transcription, and whether these factors target *dhd* directly and simultaneously, would be essential to understand the cascade of events leading to its massive expression.

The case of *dhd* indeed illustrates the complexity of understanding the chromatin landscape at cell type-specific genes, when the starting material is a complex tissue. In this context, the Cut&Run analysis implemented in our study allowed us to reveal the co-occupancy of H3K27me3 nucleosomes and associated transcription factors. While this approach cannot identify the cell of origin of each individual DNA molecule, it can be used to make important deductions on the combinatorial co-occupancy on DNA of different chromatin components. This approach joins other recent methods comparable in their principle, namely the DNA methyl-transferase single-molecule footprinting (dSMF) method [61] and the low-salt antibody-targeted tagmentation (CUTAC) approach [62]. Together with single-cell methodologies, these approaches hold the potential to begin uncovering complex epigenomic regulation processes, such as that of *dhd*, that were until recently inaccessible.

## ACKNOWLEDGEMENTS

We thank Dr. Kami Ahmad for sharing reagents and advice on Cut&Run. We are grateful to the TRiP projects and Bloomington Drosophila Stock Center for fly stocks. We acknowledge the contribution of SFR Biosciences (UAR3444/CNRS, US8/Inserm, ENS de Lyon, UCBL) facilities PLATIM and Arthro-tools. Sequencing was performed by the GenomEast platform, a member of the ‘France Génomique’ consortium. We would like to thank Raphaëlle Dubruille and Francesca Palladino for critical reading of the manuscript. We also warmly thank them, as well as Lucas Waltzer and all members of the Loppin lab for their input throughout this project. This work was supported by an Association pour la Recherche sur le Cancer (ARC) Foundation grant (PJA20191209671) to BL. DTC was supported by a PhD fellowship from the Ministère de l’Enseignement supérieur, de la Recherche et de l’Innovation and from the ARC (ARCDOC42020020001727).

## AUTHOR CONTRIBUTIONS

**Conceptualization:** Daniela Torres-Campana, Benjamin Loppin, Guillermo A. Orsi.

**Funding acquisition:** Benjamin Loppin, Guillermo A. Orsi.

**Investigation:** Daniela Torres-Campana, Béatrice Horard, Sandrine Denaud, Benjamin Loppin, Guillermo A. Orsi.

**Data curation and formal analysis:** Daniela Torres-Campana, Gérard Benoit, Guillermo A. Orsi.

**Methodology:** Daniela Torres-Campana, Guillermo A. Orsi

**Supervision:** Béatrice Horard, Benjamin Loppin, Guillermo A. Orsi.

**Validation:** Daniela Torres-Campana, Guillermo A. Orsi.

**Writing – original draft:** Daniela Torres-Campana, Guillermo A. Orsi.

**Writing – review & editing:** Daniela Torres-Campana, Béatrice Horard, Gérard Benoit, Benjamin Loppin and Guillermo A. Orsi.

## MATERIALS & METHODS

### Drosophila strains

Flies were raised at 25°C on standard medium. The following stocks were obtained from the Bloomington *Drosophila* Stock Center (simplified genotypes are given): *P{TRiP.HMS00849}attP2* (*mod(mdg4)* shRNA; #33907), *P{TRiP. HMS00363}attP2* (*Snr1* shRNA; #32372), *P{TRiP.GL00612}attP40* (*lid* shRNA; #36652), *P{TRiP.GLV21071}attP2* (*lid* shRNA; #35706), *P{TRiP.HMS00359}attP2* (*Sin3a* shRNA; #32368), *P{TRiP.HMS00066}attP2* (*E(z)* shRNA; #*33659), P{y[+t7.7]=CaryP}attP2* (Control line for TRiP RNAi lines; #B36303), *P{otu*-*GAL4::VP16.R}1; P{GAL4*-*nos.NGT}40; P{GAL4::VP16*-*nos.UTR}MVD1* (Maternal Triple Driver or “MTD-Gal4”; #31777), *P{GAL4::VP16*-*nos.UTR}MVD1* (“nos-Gal4”; #4937). Other stocks are: *w^1118^*, *Df(1)J5/FM7c* (Salz et al., 1994), *P[Mst35Ba-EGFP]* (Manier et al., 2010), *pW8-dhd^WT^* (Tirmarche et al., 2016). TRiP lines target all predicted isoforms of their respective target genes. “Control” in shRNA experiments refers to the offspring of the control line for TRiP lines crossed with the MTD-Gal4 line.

For the *pW8-dhd^ΔDRE^* mutant, two fragments were amplified by PCR from *w^1118^* genomic DNA using the primers ΔDRE-1-for/ ΔDRE-1-rev and ΔDRE-2-for/ ΔDRE-2-rev (Table S1). PCR products were assembled and cloned into the *pW8-dhd^WT^* vector (Tirmarche et al., 2016) previously digested by KpnI and BamHI using the NEBuilder HiFi DNA Assembly Cloning Kit (NEB, #E5520S). *dhd*^ΔDRE^ transgene was integrated in the *PBac{attP-3B}VK00031* platform (62E1) using PhiC31-mediated transformation (Bischof et al., 2007) and flies were generated by The Best Gene (TheBestGene.com).

### Germline knock-down and fertility tests

To obtain *KD* females, virgin shRNA transgenic females were mass crossed with transgenic Gal4 males at 25°C and females of the desired genotype were recovered in the F1 progeny. To measure fertility, virgin females of different genotypes were mated to males in a 1:1 ratio and placed for 2 days at 25°C. They were then transferred to a new vial and allowed to lay eggs for 24 hours. Embryos were counted and then let to develop for at least 36 hours at 25°C. Unhatched embryos were counted to determine hatching rates.

### Gene expression analysis by RT-QPCR

Total RNA was extracted from ovaries of 3-day-old females using the NucleoSpin RNA isolation kit (Macherey-Nagel), following the instructions of the manufacturer. 1μg of total RNA was reverse transcribed using the SuperScript^TM^ II Reverse Trancriptase kit (Invitrogen) with oligo (dT) primers. RT-qPCR reactions were performed in duplicates as described previously (Torres-Campana et al., 2020). Primer sets used are provided in Table S1.

### Immunofluorescence and imaging

Early (0–30 min) embryos laid by females of the indicated genotypes were collected on agar plates. Embryos were dechorionated in bleach, fixed in a 1:1 heptane:methanol mixture and stored at -20°C. Embryos were washed three times (10 min each) with PBS 0.1%, Triton X-100 (PBS-T) and then incubated with primary antibodies in the same buffer on a wheel overnight at 4°C. They were then washed three times (20 min each) with PBS-T. Incubations with secondary antibodies were performed identically. Embryos were mounted in DAKO mounting medium containing DAPI.

Ovaries were dissected in PBS-T and fixed at room temperature in 4% formaldehyde in PBS for 25 minutes. Immunofluorescence was performed as for embryos. Ovaries were then mounted as described above.

Antibodies used are provided in Table S2. Images were acquired on an LSM 800 confocal microscope (Carl Zeiss). Images were processed with Zen imaging software (Carl Zeiss) and ImageJ software.

### Western blotting

Ovaries from 30 females were collected and homogenized in lysis buffer (20mM Hepes pH7.9, 100mM KCl, 0.1mM EDTA, 0.1mM EGTA, 5% Glycerol, 0.05% Igepal and protease inhibitors (Roche)). Protein extracts were cleared by centrifugation and purified with Pierce™ GST Spin Purification Kit (ThermoFisher Scientific, #16106). Western analysis was performed using standard procedures and used antibodies and concentrations are presented in Table S2.

### Ovarian RNA sequencing and analysis

Samples were processed as previously described (Torres-Campana et al., 2020).

Sequencing was completed on two biological replicates of the following genotypes:

*mod(mdg4)* KD (*MTD-Gal4*>*shRNA mod(mdg4)*), i.e

*P{w[+mC] = otu-GAL4::VP16.R}1, w[*]/y*[1] *sc[*] v*[1]*; P{w[+mC] = GAL4-nos.NGT}40/+; P{w[+mC] =* GAL4::VP16-nos.UTR}CG6325[MVD1]/P{y[+t7.7] v[+t1.8] = TRiP. HMS00849} attP2

*Snr1* KD (*MTD-Gal4*>*shRNA Snr1*), i.e *P{w[+mC] = otu-GAL4::VP16.R}1, w[*]/y*[1] *sc[*] v*[1]*;P{w[+mC] = GAL4-nos.NGT}40/+; P{w [+mC] = GAL4::VP16-nos.UTR}CG6325[MVD1]/P{y[+t7.7] v[+t1.8] = TRiP. HMS00363}attP2*

### Chromatin profiling by CUT&RUN

Cut&Run in *Drosophila* tissues was previously described [37]. Briefly, ovaries from 3-day-old flies were dissected in Wash+ Buffer (20 mM HEPES pH 7.5, 150 mM NaCl, 0.9 mM spermidine, 0.1% BSA with cOmplete protease inhibitor, Roche) and were bound to BioMag Plus Concanavalin-A-conjugated magnetic beads (ConA beads, Polysciences, Inc). Tissues were then permeabilized for 10min in dbe+ Buffer (20 mM HEPES pH 7.5, 150 mM NaCl, 0.9 mM spermidine, 2 mM EDTA, 0.1% BSA, 0.05% digitonin and protease inhibitors). Samples were then incubated with gentle rocking overnight at 4°C with primary antibody solution in dbe+ buffer (see Table S2 for antibody concentrations). Protein A fused to micrococcal nuclease (p-AMNase) was added in dbe+ buffer and samples were incubated with rotation at room temperature for 1 hour. Cleavage was done in WashCa+ buffer (20 mM HEPES pH 7.5, 150 mM NaCl, 0.9 mM spermidine, 0.1% BSA, 2 mM CaCl2 with and protease inhibitors) at 0° for 30 minutes. Digestion was stopped with addition of 2XSTOP Buffer (200mM NaCl, 20mM EDTA, 4mM EGTA, 62.5µg/mL RNaseA). Samples were incubated at 37°C for 30 min to digest RNA and release DNA fragments. Cleaved DNA was then recovered with Ampure XP beads (Beckman Coulter) immediately after protease treatment. Antibodies used for CUT&RUN are given in Table S2. Retrieved DNA was used either for qPCR or for library preparation followed by deep sequencing. Sequencing libraries for each sample were synthesized using Diagenode MicroPlex Library Preparation kit according to supplier recommendations (version 2.02.15) and were sequenced on Illumina Hiseq 4000 sequencer as Paired-End 100 base reads following Illumina’s instructions (GenomEast platform, IGBM, Strasbourg, France). Image analysis and base calling were performed using RTA 2.7.7 and bcl2fastq 2.17.1.14. Adapter dimer reads were removed using DimerRemover.

### Cut&Rut-qPCR

0,1 ng of retrieved DNA in Cut&Run were used as template in a real time quantitative PCR assay using SYBR® Premix Ex TaqTM II (Tli RNaseH Plus) (Takara). All qPCR reactions were performed in duplicates using Bio-Rad CFX-96 Connect system with the following conditions: 95°C for 30s followed by 40 cycles of denaturation at 95°C for 5s and annealing and extension at 59°C for 30s. As a normalization control, we processed ovary samples from each studied genotype as for Cut&Run, except the antibody and pA-MNase incubation steps were omitted and instead we incubated tissue with 10U of Micrococcal Nucleasease for 30min at 37°C (ThermoFisher Scientific, #88216). Fold change in histone mark enrichment was determined relative to this whole MNase control and relative to the *Sas10* gene, which was depleted in the histone marks tested in this study. Primer sets used are provided in Table S1. Statistical tests were performed using GraphPad Prism version 6.00 for Mac OS X (GraphPad Software).

### Sequencing data processing

Paired-end reads were mapped to the release 6 of the *D*. *melanogaster* genome using Bowtie2 (v. 2.4.2). To compare samples with identical readcount, we employed Downsample SAM/BAM (Galaxy Version 2.18.2.1). We used bamCoverage from deepTools 2.0 (Galaxy Version 3.3.2.0.0) to calculate read coverage per 25bp bin with paired-end extension. Peak calling was done on sorted short fragments (<120 bp) with MACS2 (v. 2.1.1.20160309) with the following parameters: –nomodel, –p-value =0.0001, –keep-dup=all and the rest by default. To establish a high-confidence short fragment peak list we retained peaks that were present in biological replicates from the control genotype. Genome browser views screenshots were produced with the IGV software, for Cut&Run we used a 25bp bin for all fragments and a 10bp bin for short fragments (<120bp). For the midpoint-plot of fragment sizes around short fragment peaks, the length of each fragment was plotted as a function of the distance from the fragment midpoint to the summit of the peak identified by MACS2.

Heatmaps were generated with RStudio (RStudio Team (2016). RStudio: Integrated Development for R. RStudio, Inc., Boston, MA URL http://www.rstudio.com/) and the packages ‘gplots’ (v.3.1.1) and ‘plyr’ (Wickham, 2011).

### Motif scanning

Motif scanning on the *pW8-dhd^WT^* transgene sequence was done with FIMO (v. 5.3.3) (Grant et al., 2011) using the flyreg v.2 motif database with default parameters.

### Data Availability

Original sequencing data from this publication have been deposited to the Gene Expression Omnibus with identifiers GSE174263 (RNA-seq) and GSE174250 (Cut&Run).

REVIEWERS can access this data with the following tokens: For Cut&Run (GSE174250): uzexooqybxmjpud

For RNA-seq (GSE174263): idknsuamprevpsl

Additional sequencing data used in this study are available from GEO under the following accession numbers: GSE151981 (ATAC-seq), GSE37444 and GSE146993 (H3K27me3 ChIP-seq), GSE36393 (Mod(mdg4) ChIP-seq), GSE62904 (CP190, Beaf-32 and Dref ChIP-seq) and GSE24521 (Polycomb and Polyohomeotic ChIP-seq).

## SUPPORTING INFORMATION CAPTIONS

**Fig S1.**
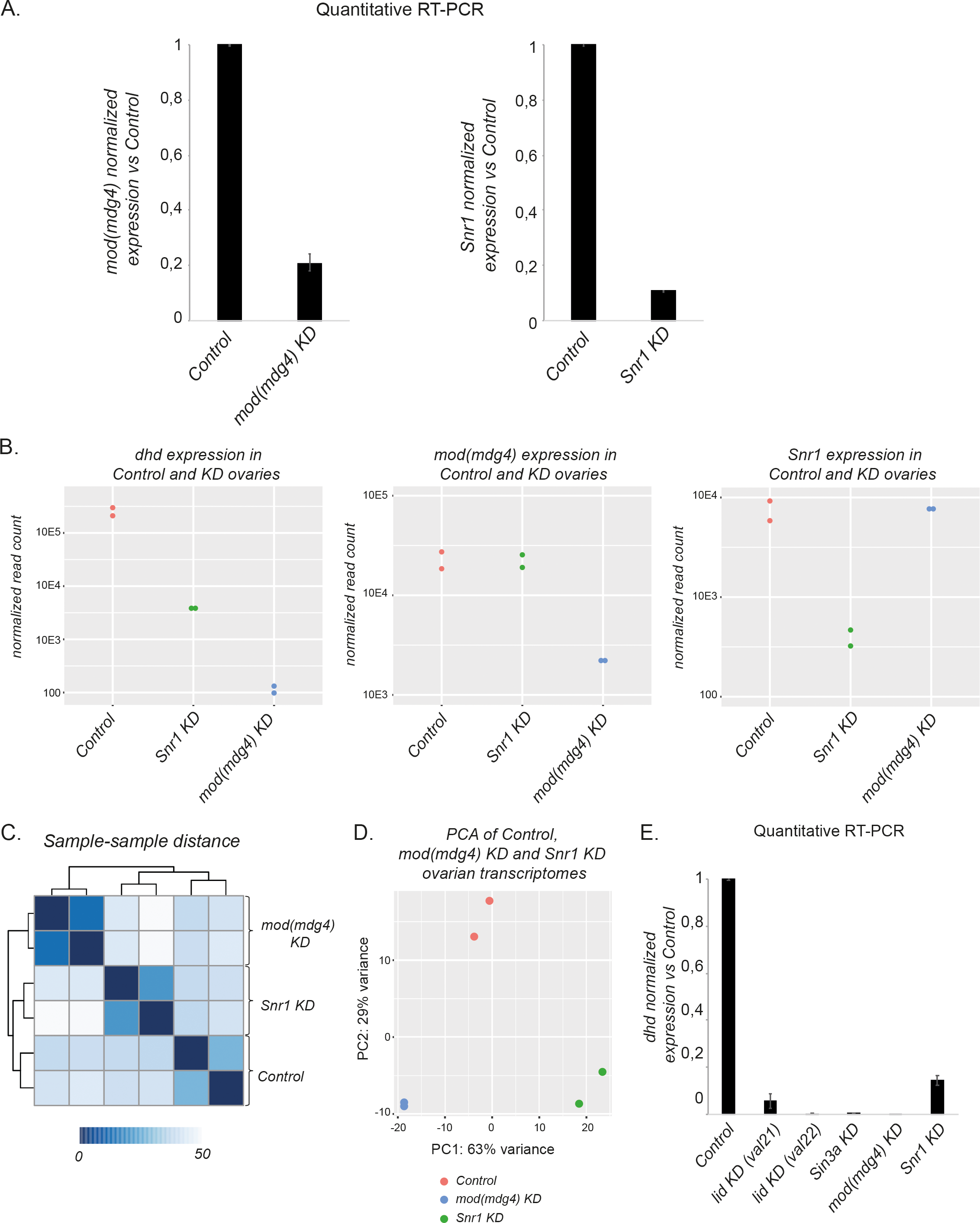
*mod(mdg4)* KD and *Snr1* KD downregulate *dhd*. A-*mod(mdg4)* KD and *Snr1* KD are efficient in the female germline. Left: RT-qPCR quantification of *mod(mdg4)* mRNA levels in Control and KD ovaries. Right: RT-qPCR quantification of *Snr1* mRNA levels in control and KD ovaries (normalized to *Rp49* and relative to expression in Control ovaries). Data from biological duplicates analyzed in technical duplicates are presented as mean ± SEM. B-Quantification of *dhd, mod(mdg4)* and *(Snr1)* counts in RNA-seq data from Fig1B. Both duplicates are shown. C-Limited overlap in the effects of *mod(mdg4)* and *Snr1* KDs. Hierarchical clustering of sample distance heatmap of RNA-seq samples. D-Principal component analysis for RNA-seq samples. E-*lid, Sin3a, mod(mdg4) and Snr1* KD severely downregulate *dhd* expression. RT-qPCR quantification of *dhd* mRNA levels in ovaries of indicated genotypes (normalized to *rp49* and relative to expression in Control ovaries). Two different shRNA constructs (val21 and val22) against *lid* were tested. Data from biological duplicates analyzed in technical duplicates are presented as mean ± SEM.

**FigS2.**
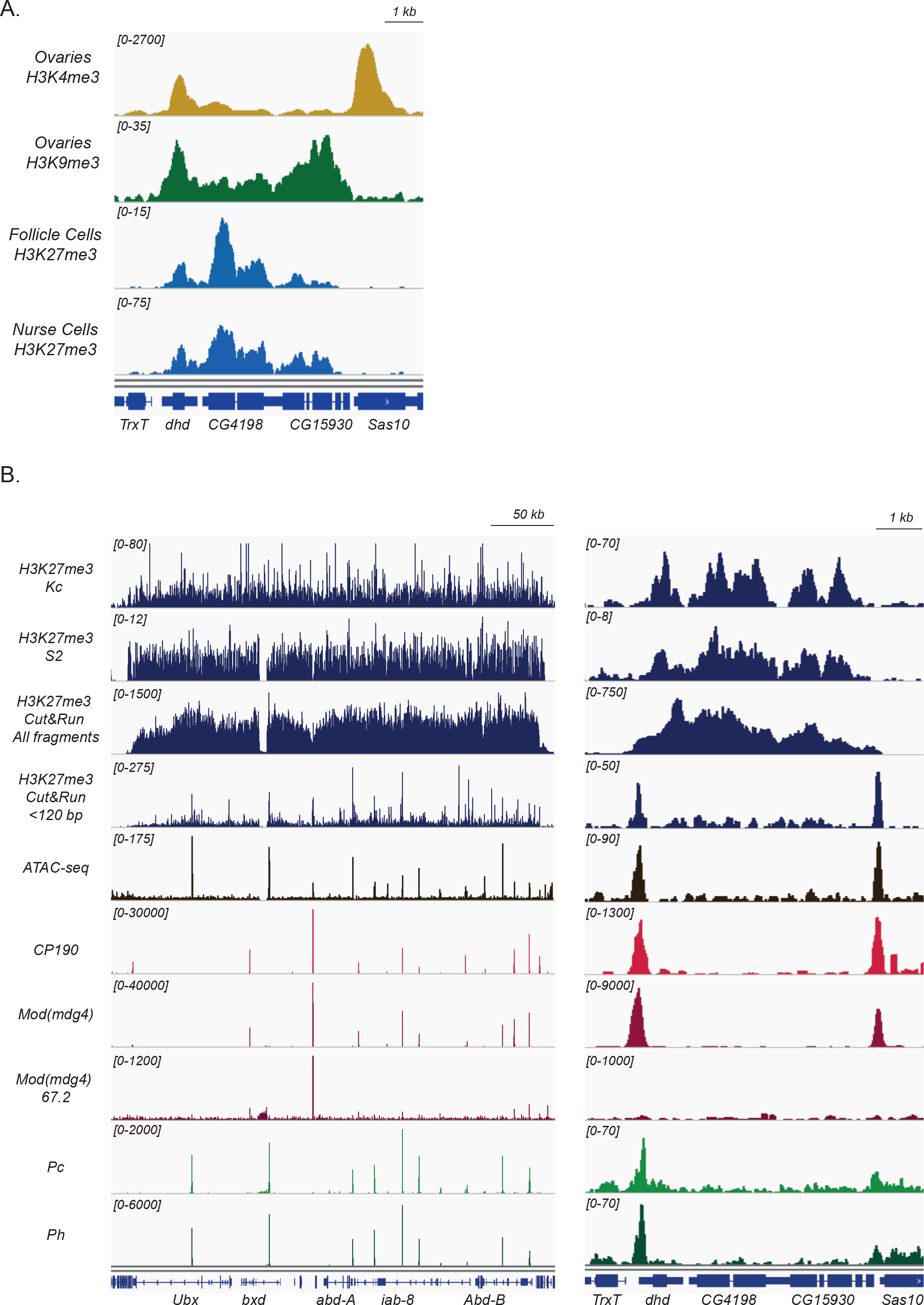
Cut&Run is consistent with ChIP-seq data. A—Histone modification profiles at the *dhd* region. ChIP-seq data showing the active mark H3K4me3 (yellow) (*Torres-Campana et al., 2020*), and the repressive marks H3K9me3 (green) (*Smolko et al., 2018*) and H3K27me3 (light blue) (*DeLuca et al., 2020*). B—Short fragment peaks align with known regulatory elements. Genome browser views of the bithorax complex (BX-C) (left) and the *dhd* region (right). Display of H3K27me3 ChIP-seq (from Kc cells, *Van Bortle et al., 2012* and S2 cells, accession number GSE146993), H3K27me3 Cut&Run (from Control ovaries, all fragments and <120bp fragments), ATAC-seq (from S2 cells, *Jain et al., 2020*), CP190 ChIP-seq (from Kc cells, *Li et al., 2015*), Mod(mdg4) (all isoforms) and Mod(mdg4)67.2 isoform ChIP-seq (from Kc cells, *Van Bortle et al., 2012*) Polycomb (Pc) and Polyhomeotic (Ph) ChIP-seq (from S2 cells, *Enderle et al., 2011*). Cut&Run short fragments largely overlap with peaks from the other tracks displayed.

**Fig S3.**
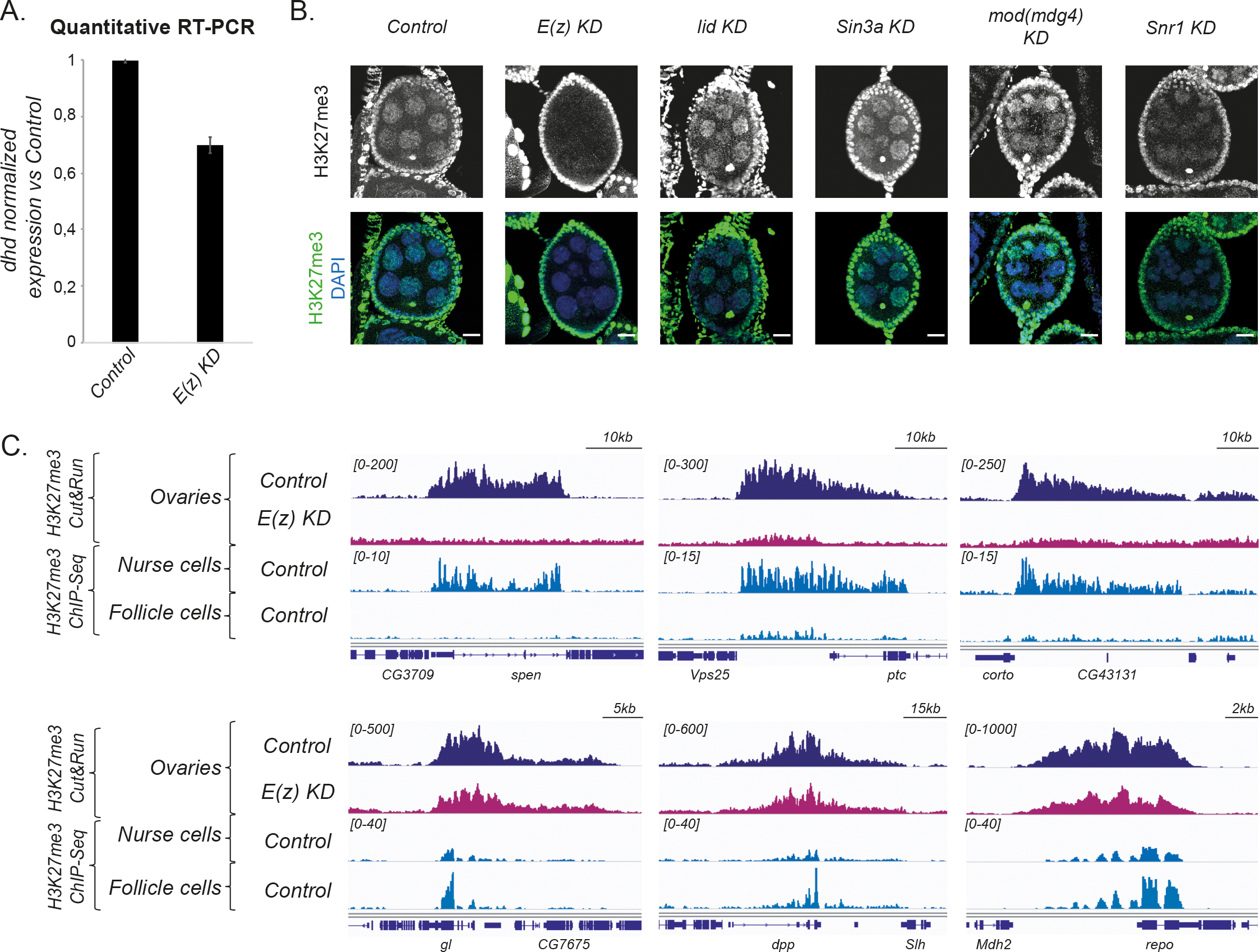
Whole-ovary experiments yield signal from both somatic follicle cells and germline cells. A-*E(z)* KD does not severely affect *dhd* expression. RT-qPCR quantification of *dhd* mRNA levels in Control and *E(z)* KD ovaries (normalized to *rp49* and relative to expression in Control ovaries). Data from biological duplicates analyzed in technical duplicates are presented as mean ± SEM. B- *E(z)* KD and *Snr1* KD affect H3K27me3 levels in nurse cells. Confocal images of representative egg chambers in Control, *E(z)* KD, *lid* KD, *Sin3a* KD, *mod(mdg4)* KD and *Snr1* KD. In control ovaries, H3K27me3 staining marks somatic follicle cell nuclei, the karyosome and germline nurse cell nuclei. In *E(z)* KD ovaries the karyosome and nurse cells loose staining of the histone mark but follicle cells are marked normally. No notable change is observed in *lid* KD, *Sin3a* KD or *mod(mdg4)* KD while in *Snr1* KD nurse cells staining is less intense. Scale bar 10μm. C- Cut&Run in whole ovaries captures signal from both somatic and germline cells. Genome browser views of H3K27me3 Cut&Run signal in Control and *E(z)* KD ovaries and H3K27me3 ChIP-seq from FACS sorted nurse cells and somatic follicle cells (*DeLuca et al., 2020*). Upper panels show representative loci enriched for the mark solely in nurse cells (germline) and absent in *E(z)* KD ovaries. Lower panels show H3K27me3 domains where the signal comes almost exclusively from follicle cells and is not significantly affected in the germline *E(z)* KD.

**Fig S4.**
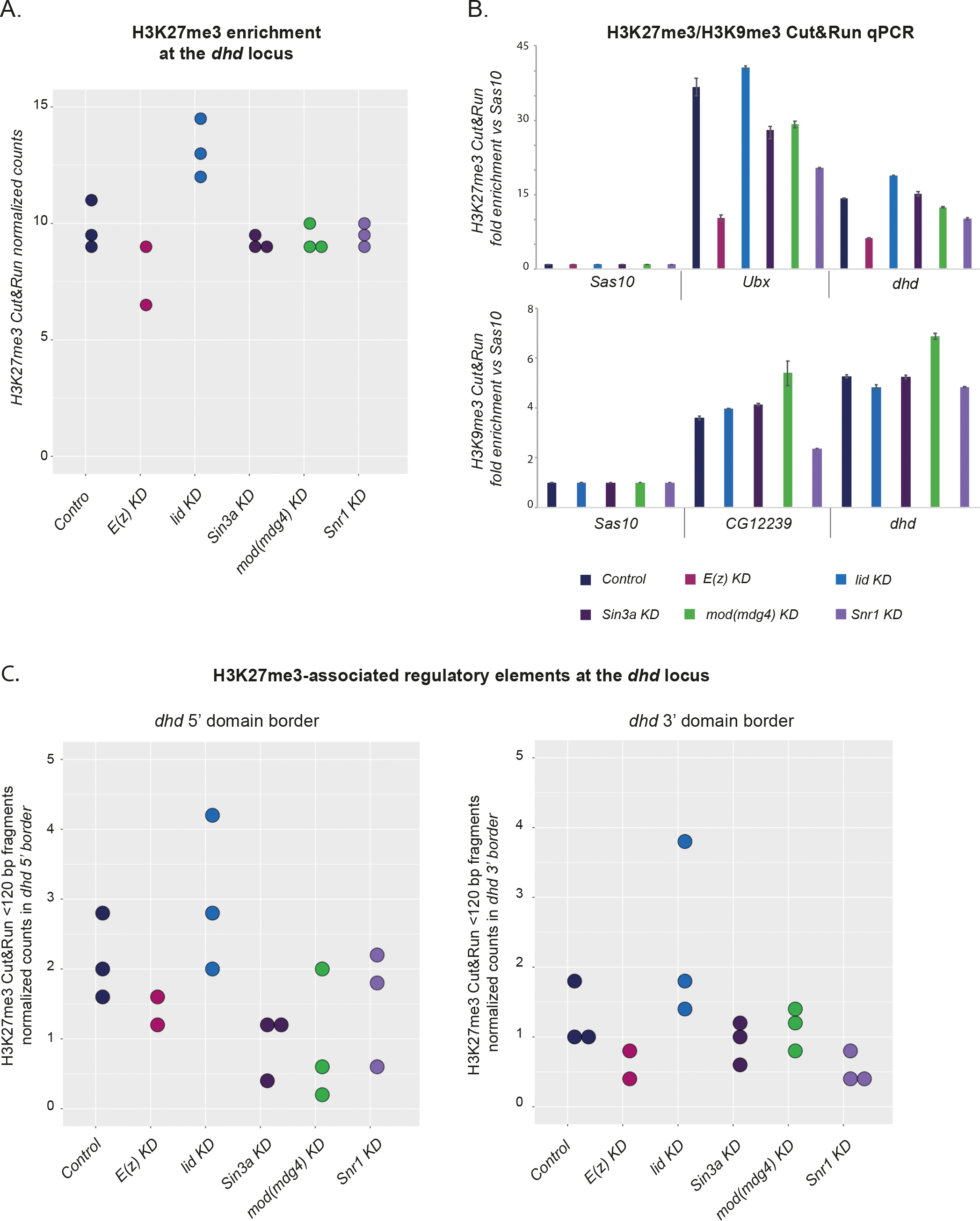
Cut&Run is reproducible among replicates and detectable by qPCR. A—H3K27me3 Cut&Run signal at the *dhd* locus from Control and KD ovaries. Left: Dotplot showing normalized read counts of H3K27me3 Cut&Run at the *dhd* domain from independent biological triplicates of the indicated genotypes (duplicates for *E(z)* KD). B—Studied KDs do not radically affect heterochromatic marks signal at *dhd*. Cut&Run-qPCR in Control and KD ovaries for H3K27me3 and H3K9me3. The *Sas10* gene was used as negative control and positive controls were *Ubx* for H3K27me3 and *CG12239* for H3K9me3. Fold enrichment was calculated relative to *Sas10.* The mean of a representative replicate analyzed in technical duplicates is shown. C— *Sin3a* KD*, Snr1* KD and *mod(mdg4)* KD affect the stability of the H3K27me3-associated regulatory elements at the *dhd* mini-domain. Dotplot showing normalized read counts of H3K27me3 Cut&Run <120bp fragments at *dhd* regulatory elements from independent biological triplicates of the indicated genotypes (duplicates for *E(z)* KD). 5’ and 3’ border elements are plotted separately.

**Fig S5.**
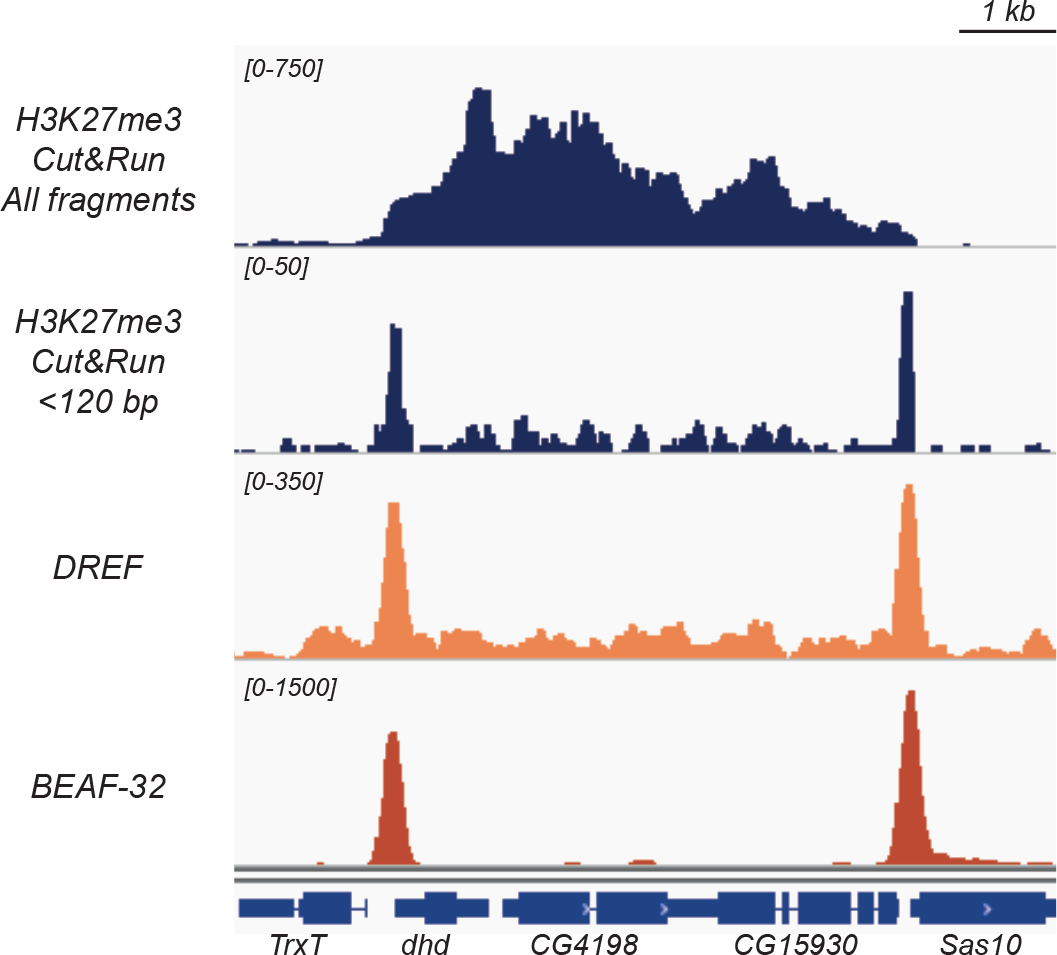
Dref and Beaf-32 are found at *dhd* regulatory elements. Genome browser views of ovarian H3K27me3 Cut&Run (all fragments and <120bp fragments) and Dref and Beaf-32 ChIP-seq (from Kc cells, *Li et al., 2015*). <120 bp fragment peaks at the *dhd* domain borders align with DREF and Beaf-32 peaks.

**Fig S6.**
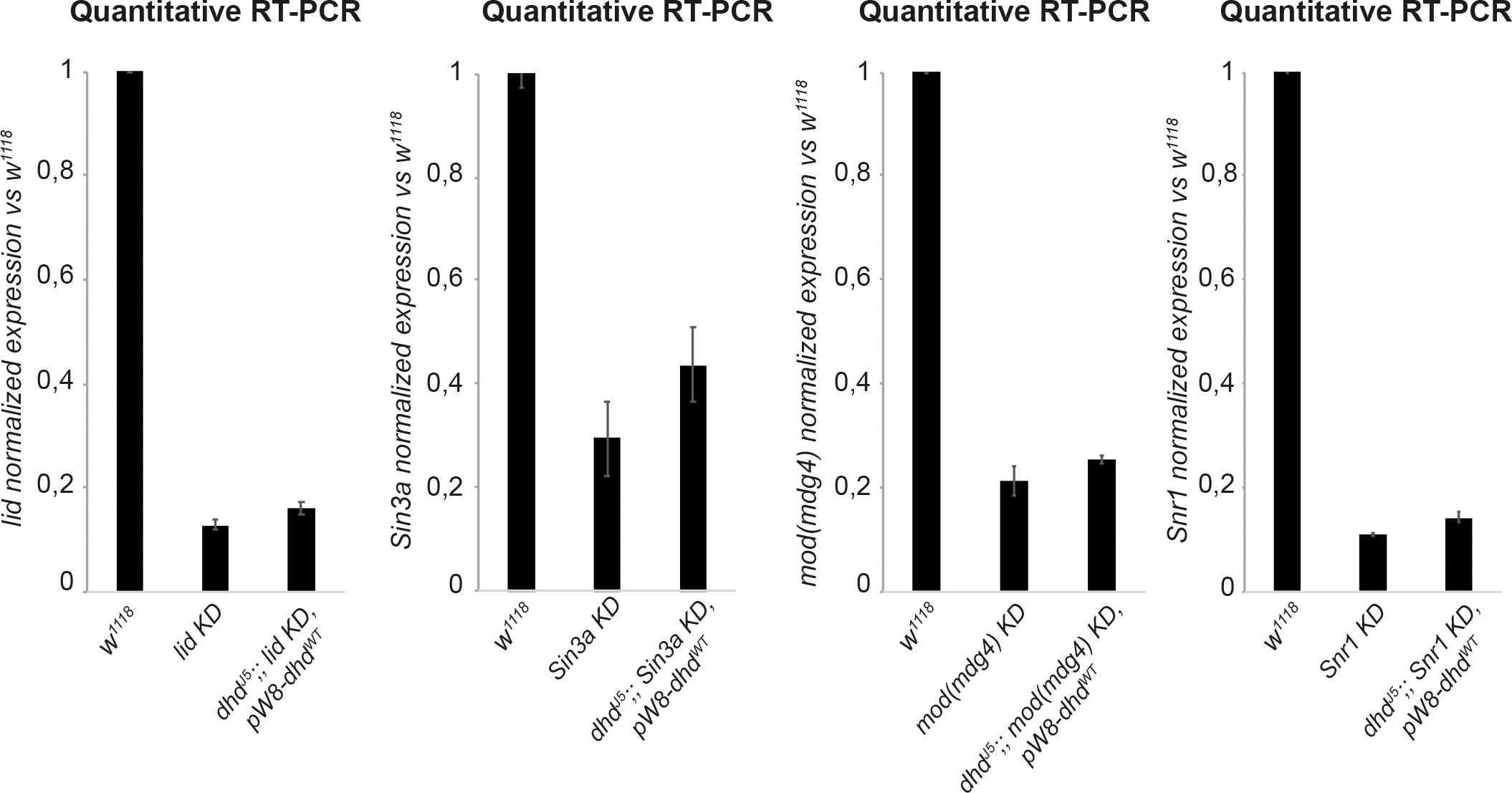
*Sin3a* KD, *mod(mdg4)* KD and *Snr1* KD are efficient in the female germline of rescue flies. From left to right: RT-qPCR quantification of *lid*, *Sin3a*, *mod(mdg4)* and *Snr1* mRNA levels in ovaries of the indicated genotypes (normalized to *rp49* and relative to expression in *w^1118^* ovaries). Data from biological duplicates analyzed in technical duplicates are presented as mean ± SEM.

**Table S1.**
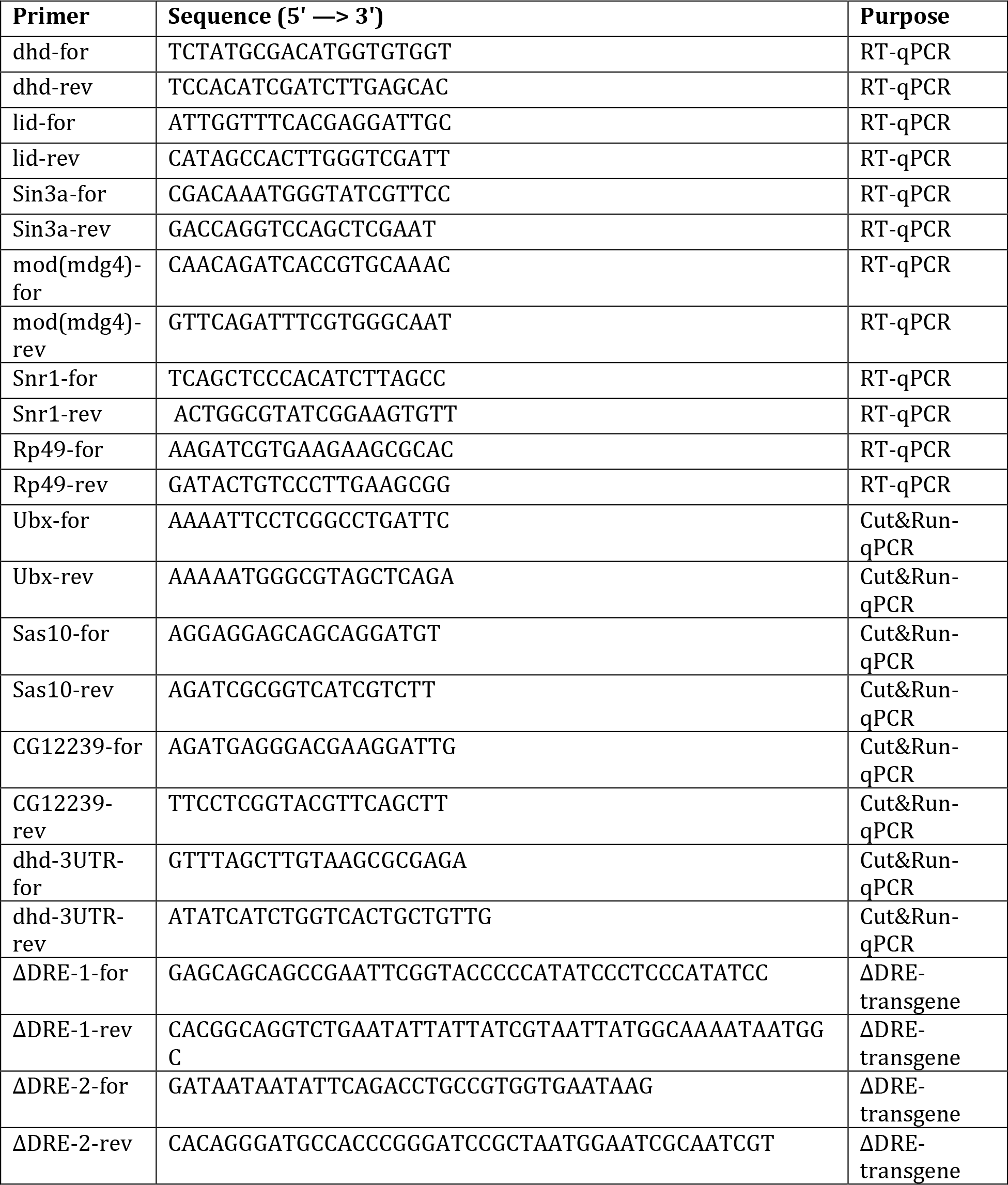
List of primers used in this paper.

**Table S2.** List of antibodies used in this paper.

| **Antibody** | **Host animal** | **Dilution** | **Experiment** | **Company (Catalog#)** |
| --- | --- | --- | --- | --- |
| Anti-Histones | mouse,<br>monoclonal | 1:1000 | Immunofluorescence | Merck (#F152.C25.WJJ) |
| Anti-GFP | mouse,<br>monoclonal | 1:200 | Immunofluorescence | Roche (#118144600001) |
| Anti-H3K27me3 | rabbit,<br>polyclonal | 1:500 | Immunofluorescence | Merck (#07-449) |
| Anti-H3K27me3 | rabbit,<br>monoclonal | 1:100 | Cut&Run | Cell Signalling Technology<br>(#9733) |
| Anti-H3K9me3 | rabbit,<br>polyclonal | 1:50 | Cut&Run | Abcam (#8898) |
| Anti-DHD | rabbit,<br>polyclonal | 1:1000 | Western Blot | *Tirmarche et al., 2016* |
| Anti-α-tubulin | mouse,<br>monoclonal | 1:500 | Western Blot | Merck (#T9026) |

